# Associative Learning and Motivation Differentially Requires Neuroligin-1 at Excitatory Synapses

**DOI:** 10.1101/2020.01.01.890798

**Authors:** Jiaqi Luo, Jessica M Tan, Jess Nithianantharajah

**Affiliations:** Florey Institute of Neuroscience and Mental Health, 30 Royal Parade, Parkville Victoria Australia; Florey Department of Neuroscience, University of Melbourne, Parkville Victoria, Australia

## Abstract

In a changing environment, a challenge for the brain is to flexibly guide adaptive behavior towards survival. Understanding how these decision-making processes and underlying neural computations are orchestrated by the structural components of the brain, from circuits to cells, and ultimately the signaling complex of proteins at synapses, is central to elucidating the mechanisms that shape normal and abnormal brain connectivity, plasticity and behavior. At excitatory synapses, neuroligin-1 (Nlgn1) a postsynaptic cell-adhesion molecule required for the formation of trans-synaptic complexes with presynaptic partners is critical for regulating synapse specification, function and plasticity. Extensive evidence shows Nlgn1 is essential for synaptic transmission and long-term plasticity, but how these signaling processes ultimately regulate components of cognitive behavior is much less understood. Here, employing a comprehensive battery of touchscreen-based cognitive assays, we measured two key decision problems: i) the ability to learn and exploit the associative structure of the environment and ii) the trade-off between potential rewards and costs, or positive and negative utilities associated with available actions. We found that mice lacking Nlgn1 have an intact capacity to acquire complex associative structures and adjust learned associations. However, loss of Nlgn1 alters motivation leading to a *reduced* willingness to overcome response effort for reward and an *increased* willingness to exert effort to escape an aversive situation. We suggest Nlgn1 may be important for balancing the weighting on positive and negative utilities in reward-cost trade-off. Our findings identify Nlgn1 is essential for regulating distinct cognitive processes underlying decision-making, providing evidence of a new model for dissociating the computations underlying learning and motivational processing.

## INTRODUCTION

In a changing environment, a challenge for the brain is to adaptively guide behavior towards survival which involves the processing of sensory information, selecting between actions that will most likely result in a beneficial outcome, and executing these actions. The foundation of these cognitive capacities is hardwired in the architecture of the brain (1). Understanding how adaptive behavioral processes are orchestrated by the structural components of the brain, from circuits to cells and ultimately the signaling complex machinery of proteins at synapses, is central to elucidating the mechanisms that shape normal and abnormal brain connectivity, plasticity and behavior. At the postsynaptic density (PSD) of excitatory synapses, neuroligin-1 (Nlgn1) is a cell adhesion molecule that is critical for synapse specification, function and plasticity. It binds presynaptic neurexins to form trans-synaptic complexes (2, 3). Aligning PSD components with presynaptic neurotransmitter release sites (4–6), Nlgn1 directly binds postsynaptic scaffolds including PSD-95 (4), promotes retention of α-amino-3-hydroxyl-5-methyl-4-isoxazole-propionate (AMPA) and *N*-methyl-D-aspartate (NMDA) receptors by indirect intracellular and direct extracellular interactions in developing and mature synapses (7–10).

Functionally, NMDA receptor-mediated postsynaptic currents have been consistently shown to be decreased across several brain regions by Nlgn1 knockout or knockdown (8, 11–20) and conversely, increased by Nlgn1 overexpression (19–21). Further, Nlgn1 has been robustly shown to be essential for synaptic plasticity and the induction of both NMDA receptor-dependent and independent long-term potentiation (LTP) in multiple brain regions (8, 11, 12, 14–17, 22, 23). Contrary to the extensive molecular and physiological characterization, fewer studies have examined Nlgn1 in behavior, and none have evaluated different components of complex cognitive behavior in detail. Blundell and colleagues reported numerous behavioral measures were unaffected in mice lacking Nlgn1, but grooming behavior and spatial memory in the Morris Water Maze was disrupted (11). Others have also reported altered spatial learning and memory in the Water Maze in Nlgn1 overexpression mouse models (21, 24) and impaired contextual and cued fear memory recall in rats with lateral amygdala knockdown of Nlgn1 (16), collectively supporting the idea that Nlgn1 is important for learning and memory.

In this study, employing a comprehensive battery of touchscreen-based cognitive assays we assessed male and female mice lacking Nlgn1 to measure the ability to solve two key decision problems: 1) the ability to learn and exploit the associative structure of the environment and 2) balancing the trade-off between potential rewards and costs, or positive and negative utilities associated with available actions. We found that mice lacking Nlgn1 have an intact capacity to acquire complex associative structures and adjust learned associations. However, loss of Nlgn1 alters motivation leading to a *reduced* willingness to overcome response effort for reward and *increased* willingness to exert effort to escape an aversive situation. We suggest these divergent phenotypes may converge on a model of increased weighting on negative utilities, highlighting a novel valence-dependent role of Nlgn1 in balancing the weighting on positive and negative utilities in reward-cost trade-off. Our findings identify Nlgn1 is essential for regulating distinct cognitive processes underlying decision-making, providing evidence of a new model for dissociating the computations underlying learning and motivational processing. Furthermore, our work contributes towards unravelling the genetic architecture of dissociable cognitive modules.

## RESULTS

### Nlgn1 is not essential for learning complex associative structures

Intact long-term forms of plasticity and NMDAR function are both synaptic mechanisms thought to be required for learning and memory (e.g., (25–33). Based on the established impairments in NMDAR function and LTP combined with the previous behavioral reports, we hypothesized that Nlgn1 is likely to be important for acquiring associative structures of the environment, and using these structures to optimize action selection. To address this, we assessed male and female null mutant mice lacking Nlgn1 (*Nlgn1*^-/-^) and control wildtype (WT) littermates in a series of rodent touchscreen cognitive tests, where mice were required to make responses via nose-pokes to different visual stimuli displayed on a touchscreen to obtain rewards under different test situations. Of note, across all the cognitive tests and analyses performed, we observed no significant interactions between genotype and sex (except for that presented in Figure 6E, data in Supplemental Figure 11), therefore combined data for both sexes by genotype will be presented for clarity. Additionally, we employed trial-by-trial analyses that better describes complex behavioral data (see Methods and Supplemental Table 1 for detailed statistical results).

**Figure 1:**
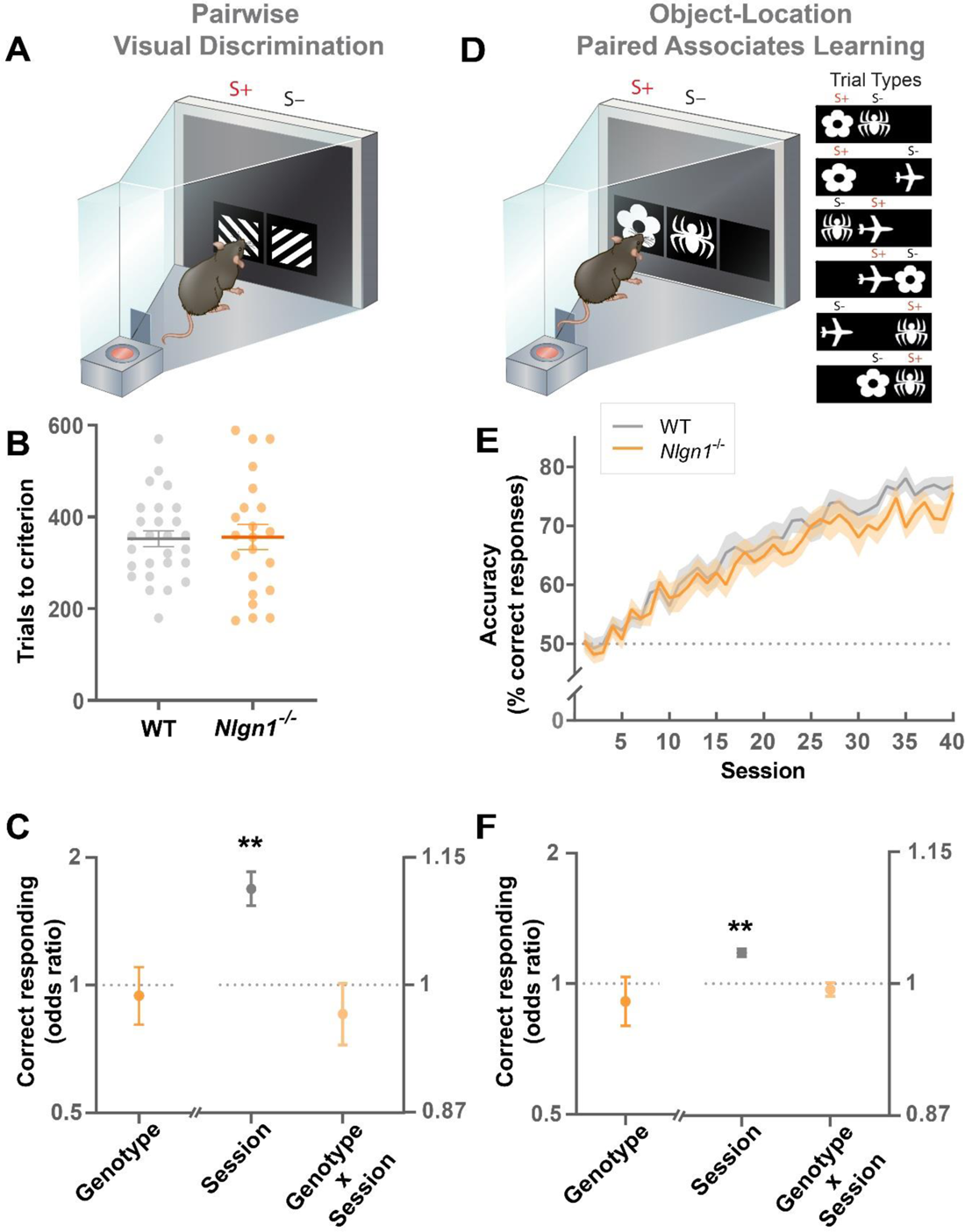
***Nlgn1*^-/-^ mice display normal associative learning**. **(A-C)** Pairwise Visual Discrimination Learning Task. **(A)** Stimuli used, S+ (rewarded stimulus), S-(unrewarded stimulus). **(B)** Number of trials to reach learning criterion; two-way ANOVA, values represent means ± SEM. **(D-F)** Object-Location Paired Associates Learning Task. **(D)** Visual stimuli and their paired correct locations (6 possible trial types). **(E)** Visuospatial learning curve showing performance accuracy across sessions, values represent means ± SEM. **(C,F)** Effect of genotype, session and their interaction on correct responding, GLLAMM (logistic link function), \*\**p* < 0.005 significantly different from 1, values represent the estimated effect of the variables on the odds of correct responding (odds ratio) ± 95% CI.

Animals first underwent several phases of instrumental training (touchscreen pre-training) to learn to initiate the commencement of trails and selectively nose-poke simple visual stimuli displayed on the touchscreen in order to obtain a liquid reward (strawberry milk) (34, 35). *Nlgn1*^-/-^ and WT mice required similar numbers of sessions to complete the pre-training phases, indicating loss of Nlgn1 does not impact the acquisition of simple instrumental conditioning (Supplemental Figure 1). Following pre-training, mice were introduced to the pairwise visual discrimination task, a forced choice paradigm where two visually similar stimuli were presented pseudorandomly between two locations (left or right side of the touchscreen). Responses to one of the stimuli were rewarded (S+) while responses to the other were unrewarded (S-) (Figure 1A). The visual discrimination task therefore requires mice to learn to perceptually discriminate the stimuli and selectively respond to the correct or rewarded stimulus regardless of stimulus location. Performance accuracy (% of correct responses) was the primary measure used to track learning across training sessions, until a learning criterion was reached. We found that *Nlgn1*^-/-^ mice required similar numbers of trials to reach the discrimination learning criterion as WT controls (Figure 1B). To understand how key variables including genotype, sex and session affect response accuracy, we estimated the effect of these variables on trial outcomes (correct/incorrect responses) using a mixed-effect generalized linear model (36). We observed a highly significant effect of session as expected, reflecting the improvements on response accuracy over sessions. However, there was no effect of genotype on trial outcomes (correct responding of *Nlgn1*^-/-^ relative to WT expressed as odds ratio, Figure 1C) nor a significant genotype x session interaction, indicating that both *Nlgn1*^-/-^ and WT mice acquired visual discrimination learning at a similar rate (Figure 1C), consistent with our trials to learning criterion analysis.

To increase demands on associative complexity by having to integrate both visual and spatial information as features defining the reward contingencies, we next employed the object-location paired associates learning task which requires mice to not only learn the perceptual features of three stimuli (flower, plane, spider) but also their unique rewarded location on the touchscreen (left, center, right respectively). On each trial, only two stimuli are presented: one displayed in its correct location (S+) and the other in an incorrect location (S-) (Figure 1D). Compared to the visual discrimination task, the greater difficulty in acquiring visuospatial associations in this object-location paired associates learning task is reflected in a slower rate of learning, evident by the increased number of training sessions (Figure 1E) and smaller effect size of session (Figure 1F). Despite this increased difficulty, we again observed *Nlgn1*^-/-^ mice were able to acquire the complex object-location associations similar to WT mice (Figure 1E-F).

### Flexible updating of learned associations is not altered by loss of Nlgn1

Adapting to dynamic environments where the outcome of a response is not always stable requires the ability to inhibit learned responses once they no longer yield positive outcomes and explore alternatives. To examine the requirement of Nlgn1 for flexible adjustment of response selection, we employed two tests. First, we examined cognitive flexibility in a test of reversal learning. Once mice had reached the learning criterion on pairwise visual discrimination (Figure 1A-C), we reversed the reward contingencies so that the previously rewarded stimulus was now unrewarded and vice versa (Figure 2A). All mice displayed a strong tendency to respond to the previously rewarded stimulus (now S-) at the beginning of reversal learning, then gradually shifted their responding to the updated S+ (previous S-) as expected (Figure 2B). We observed no differences in response accuracy between *Nlgn1*^-/-^ and WT mice throughout reversal learning (effect of genotype), nor the rate of reversal learning (genotype x session interaction) (Figure 2C).

**Figure 2:**
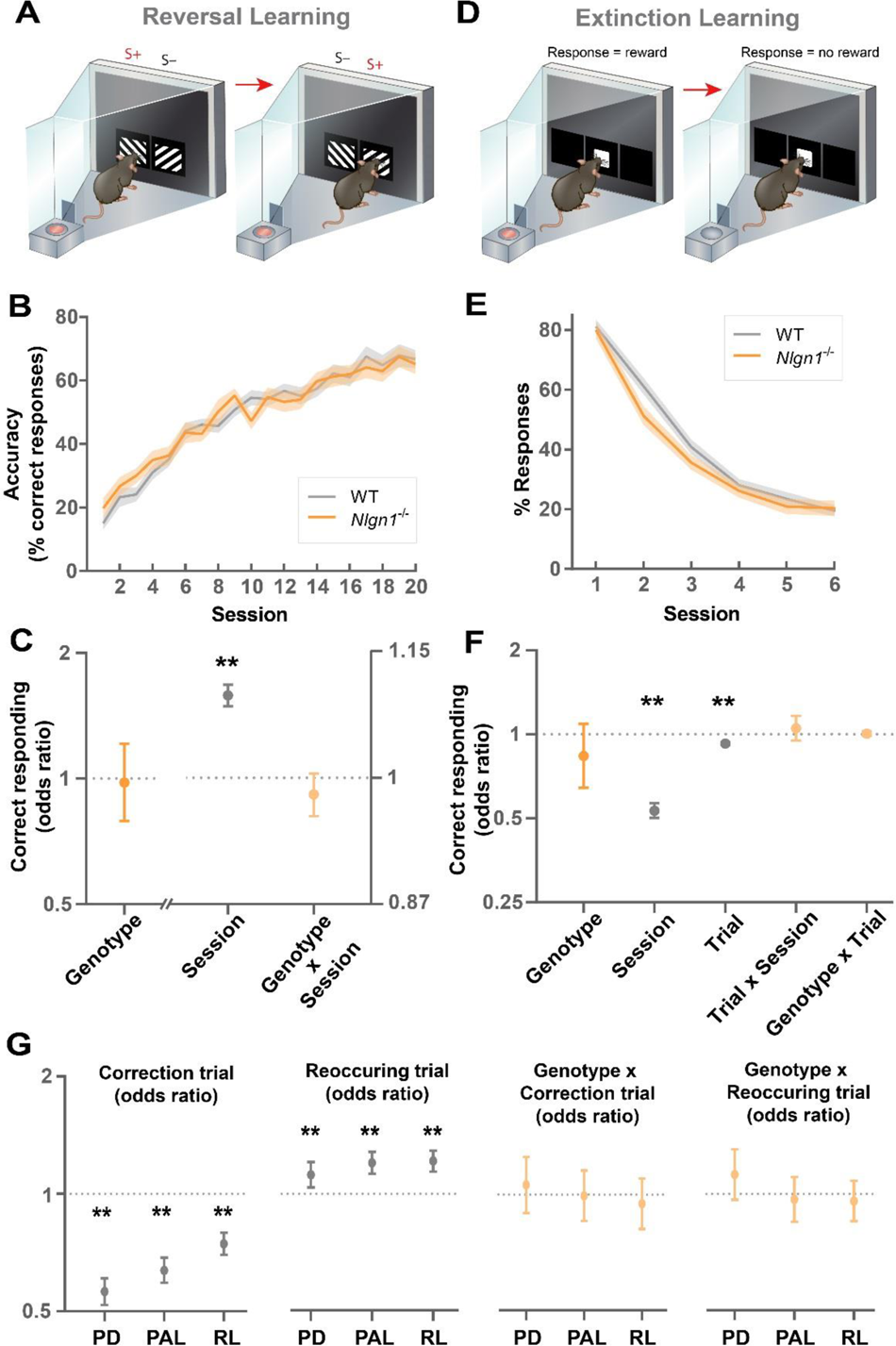
***Nlgn1*^-/-^ mice display normal adjusting of learned associations**. **(A-C)** Reversal Learning Task. **A)** Reward contingencies were switched following acquisition of learning criterion for visual discrimination, S+ (rewarded stimulus), S-(unrewarded stimulus). **(B)** Reversal learning curve showing performance accuracy across sessions, values represent means ± SEM. **(C)** Effect of genotype, session and their interaction on correct responding, GLLAMM (logistic link function), \*\**p* < 0.005 significantly different from 1, values represent odds ratio ± 95% CI. **(D-F)** Extinction Learning Task. **(D)** Once robust instrumental responding to a performance criterion was reached, responses were no longer rewarded. **(E)** Extinction learning curve showing percentages of responses across sessions, values represent means ± SEM. **(F)** Effect of genotype, session, trial within a session and their interactions on responding, GLLAMM (logistic link function), \*\**p* < 0.005 significantly different from 1, values represent the estimated effect of the variables on the odds of correct responding (odds ratio) ± 95% CI. **(G)** *Nlgn1*^-/-^ and WT mice both display similar levels of perseverative behavior in response selection across tasks (Pairwise Visual Discrimination, PD; Object-Location Paired Associates Learning, PAL; Reversal Learning, RL). Mice were less accurate on correction trials (effect of correction trial on correct responding, effect sizes < 1); more accurate on pseudorandom trials with reoccurring stimulus configurations (effect of reoccurring pseudorandom trial on correct responding, effect size > 1); but there were not differences due to genotype (effect of correction trial x genotype interaction, effect of reoccurring pseudorandom trial x genotype interaction). GLLAMM (logistic link function), \*\**p* < 0.005, values represent the estimated effect of the variables on the odds of correct responding (odds ratio) ± 95% CI.

Second, we investigated extinction learning measuring the rate at which mice stop making a learned instrumental response when those responses no longer result in an outcome, and there are no competing operant response alternatives. Mice were first trained to robustly respond to a simple stimulus (white square) and once a stable performance criterion was reached, extinction was tested in which responses to the stimulus were no longer rewarded (Figure 2D). On each trial, mice could either make or omit a response within a set time window. One challenge in quantifying the rate of instrumental extinction is that feedback on reward contingency is only provided after a response. As a result, animals that display slower instrumental extinction for example (therefore made more responses) will have received more learning opportunities due to greater feedback, and vice versa. To minimize potential differences in learning opportunities, we set a limit on the number of trials per session for all animals and assessed extinction learning across sessions (Figure 2E, Supplemental Figure 2). We observed no difference between *Nlgn1*^-/-^ and WT mice on whether they responded or omitted a trial throughout extinction (effect of genotype), rates of extinction both within a session (effect of trial, genotype x trial) and across sessions (effect of session, genotype x session) (Figure 2F). To confirm potential differences in extinction rate was not masked by differences in learning opportunities, we also analyzed the rate of extinction as the effect of cumulative responses and again found no differences (data not shown) indicating intact instrumental extinction learning.

Previous work has reported that *Nlgn1*^-/-^ mice display increased self-grooming (11) thought to represent repetitive and stereotypic behavior common in brain disorders such as autism spectrum disorder and obsessive compulsive disorder (37, 38). We were therefore interested to explore measures of repetitive or perseverative response selection across our multiple learning assays. In the pairwise visual discrimination, object-location paired associates learning and reversal learning tasks, a correct response to a first-presentation pseudorandom trial (referred to as a ‘trial’) was always followed by another trial, where the stimuli and location are displayed in a pseudorandom and counterbalanced manner. In contrast, an incorrect response was always followed by a ‘correction trial’ where the exact same stimulus-location configuration of that (pseudorandom) trial is repeatedly presented until mice switch their response to make a correct response and earn a reward (Supplemental Figure 3A). A perseveration index (PI) calculated as the average number of correction trials committed per incorrect response has been commonly used to measure perseverative responding (e.g., (33, 34, 39). However, to more explicitly and quantitatively examine perseverative responding on correction trials, we estimated the effect of correction trials on correct responding. Mice were less accurate on correction trials suggesting a tendency to reselect the same incorrect response previously selected (Figure 2G). Consistent with a tendency of repeating the previous response, mice are more accurate when the same stimulus-location configuration happened to reoccur on a consecutive trial (referred to as a ‘reoccurring trial’) therefore more likely to reselect a correct response previously selected (Figure 2G). However, we found no differences in perseverative responding between *Nlgn1*^-/-^ and WT mice (genotype x correction trial, genotype x reoccurring trial) (Figure 2G). These data show mice appear to generally have a tendency towards perseverative action selection.

### Mice lacking Nlgn1 take longer to perform instrumental actions for rewards

Decision-making in the natural world involves more than choosing the response with the highest expected rewards. Actions may have very different effort requirements, therefore balancing the trade-off between rewards and costs is a crucial part of maximizing the net utility of actions. We have so far examined task measures that involve action selection between two alternatives that require the same amount of physical effort and only differ in the expected rewards (e.g., performance accuracy calculated on correct vs incorrect responding). But like most naturalistic decision problems, these touchscreen-based tasks are free-operant tasks in that mice are free to choose a wide range of other actions (e.g., exploring, resting) instead of choosing to execute actions towards earning a reward (initiating a trial, making a response, collecting a reward). To capture an animal’s engagement in performing instrumental actions in our tasks, we analyzed several latency parameters: trial initiation, response and reward collection (Figure 3A, Supplemental Figure 3B). Response latency was further separated into i) stimulus-approach latency: time taken after initiating a trial to reach the front of the chamber near the touchscreen) and ii) stimulus-selection latency: time taken from reaching the front of the chamber and making a nose-poke response to a stimuli. Dissecting the response latencies revealed stimulus-selection latencies positively predicted performance accuracy suggesting it influences the decision between correct and incorrect responses, whereas stimulus-approach latencies did not, suggesting it reflects the decision between choosing to make a response or not (Supplemental Figure 4). Analysis across all three of ours tasks (pairwise visual discrimination, object-location paired associates learning, reversal learning) consistently revealed the same pattern for the various latency measures (Figure 3). *Nlgn1*^-/-^ mice were significantly slower to initiate trials, approach stimuli and collect rewards (Figure 3B-D, Supplemental Figure 5, confirmed with quantile regressions to analyze distribution-wide differences Supplemental Figure 6). However, stimulus-selection latencies were nearly identical between *Nlgn1*^-/-^ and WT mice (Figure 3E).

**Figure 3:**
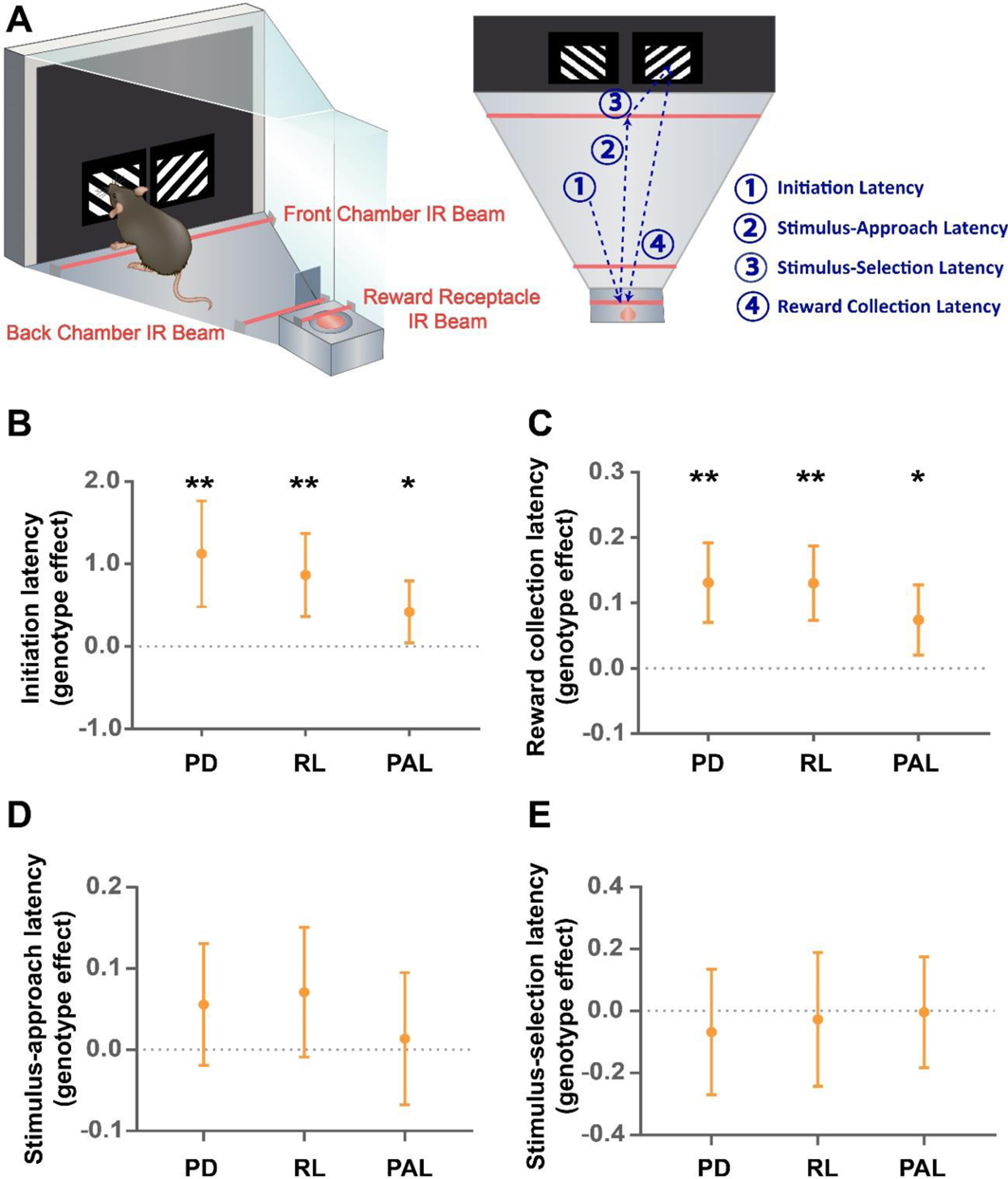
***Nlgn1*^-/-^ take longer to perform instrumental actions for rewards. (A)** Infrared-red (IR) beams located within the chambers at the front, back and reward receptacle allow the dissection of multiple reaction times (initiation latency; stimulus-approach and stimulus-selection latency; reward collection latency). *Nlgn1*^-/-^ mice took longer to **(B)** initiate trials, **(C)** collect rewards and **(D)** approach the touchscreen (stimulus-approach latency) but not **(E)** stimulus-selection across tasks (effect of genotype > 0). Pairwise Visual Discrimination (PD), Reversal Learning (RL), Object-Location Paired Associates Learning (PAL), data arranged in order of task training. Quantile regression (median), \**p* < 0.05, \*\**p* < 0.005, values represent estimated latency difference between *Nlgn1*^-/-^ and WT mice ± 95% CI.

### Loss of Nlgn1 reduces motivational processing to overcome response effort for rewards

Based on the increased latencies *Nlgn1*^-/-^ mice displayed, we hypothesized Nlgn1 may be important for regulating specific components of motivational processing. To examine this, we first tested naive mice sequentially across sessions that required a fixed ratio of responding for rewards with increasing demands (FR1-40). We wondered whether loss of Nlgn1 might reduce the number of responses at higher ratio requirements where responding has a lower utility, resembling a well-characterized model of amotivation (40–42). Indeed, when the response-reward ratio was low (FR1), *Nlgn1*^-/-^ mice performed like WT controls indicating similar motivation to respond when response utility is high and similar rate to reach satiety. However, as this ratio increased (FR5, FR20, FR40), Nlgn*1*^-/-^ mice started to make significantly fewer responses, with the reduction being greater at the higher ratio requirements (Figure 4A-B, Supplemental Figure 7). This increasing difference in responses between genotypes across ratio requirements was primarily driven by the non-linear increase in the latency taken to re-engage in responding after consuming a reward (post-reinforcement pause, Figure 4C, Supplemental Figure 8A), and the average time interval between each subsequent response (Figure 4D, Supplemental Figure 8B, Supplemental Figure 9). Next, using a separate naive cohort of mice, we wanted to see if we could observe the same motivational phenotype in a progressive ratio task where ratio requirements progressively increased within a session until mice stop responding. Indeed, we were able to reproduce the same finding with *Nlgn1*^-/-^ mice making fewer responses and therefore having a lower breakpoint relative to controls (Supplemental Figure 10).

**Figure 4:**
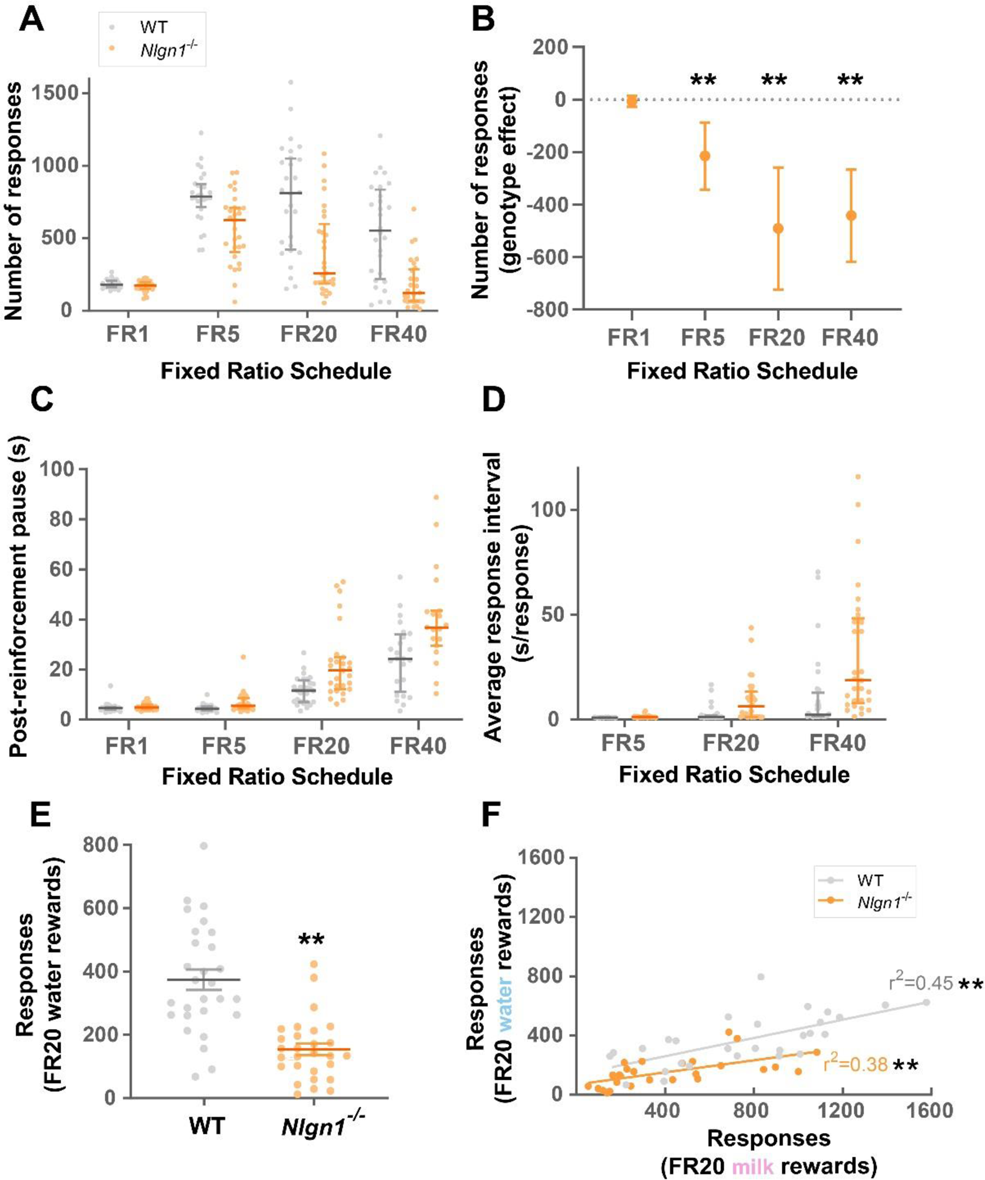
*Nlgn1*^-/-^ mice show reduced motivation to overcome response effort for rewards. **(A)** Total number of responses averaged over three sessions per ratio requirement. Bars indicate 1^st^, 2^nd^ (median) and 3^rd^ quartiles. **(B)** *Nlgn1*^-/-^ mice made fewer responses in the higher ratio requirements (effect of genotype < 0), quantile regression (median), \*\**p* < 0.005, values represent estimated effect of genotype on response counts ± 95% CI. **(C)** Post-reinforcement pause (time to the first response after consuming a reward, seconds) averaged over three sessions per ratio requirement. **(D)** Average response interval (time spent per response after an animal made the first response of a trial, seconds per response), averaged across three sessions per ratio requirement. See Supplemental Figure 9 for response-by-response breakdown of inter-response intervals. **(E)** *Nlgn1*^-/-^ mice also made fewer responses for water rewards on fixed ratio 20 (FR20), linear regression, \*\**p* < 0.005 values represent means ± SEM. **(F)** Number of responses made for milk rewards positively correlated with number of responses made for water rewards on FR20 for both WT and *Nlgn1*^-/-^ mice, linear regression, \*\**p* < 0.005.

Reduced motivation could be due to taste insensitivity to palatable rewards (e.g., strawberry milk in this case). To examine whether the observed phenotype in *Nlgn1*^-/-^ mice was specific to high-fat-high-sugar rewards, we next used water as the reward and measured responding at a fixed ratio where a robust difference was observed using strawberry milk (FR20). Similar to that observed with strawberry milk rewards, *Nlgn1*^-/-^ mice made significantly fewer responses (Figure 4E), with this effect being stronger in females compared to males (Supplemental Figure 11A). Importantly, there were significant positive correlations between the numbers of responses for strawberry milk and water rewards, indicating that mice (either WT or *Nlgn1*^-/-^) that was more motivated by strawberry milk also responded more for water (Figure 4F, Supplemental Figure 11B). Together, these data show lack of Nlgn1 impacts instrumental responding motivated by both hunger and thirst.

### *Nlgn1*^-/-^ mice exert *less* effort to earn rewards but *more* effort to escape from aversion

To investigate the generalizability of the observed motivational phenotype, we wanted to next examine whether *Nlgn1*^-/-^ mice were also less willing to exert effort outside an operant environment. We assessed exploration and spontaneous locomotor activity in a novel, open-field environment (in darkness to minimize stress) to measure the decision between exploration and resting. We tested two separate cohorts of animals, one group that had previously been tested in an operant paradigm and a second experimentally naive group. Interestingly, in the group that had previous experimental experience, we see that *Nlgn1*^-/-^ mice travel a shorter distance (Figure 5A) and spent more time resting (Figure 5B) compared to WT controls, consistent with a reduced willingness to overcome the cost of physical effort. We noted that when we assessed the naive cohort of animals, we didn’t see a significant effect of genotype (Supplemental Figure 12) suggesting these measures of exploratory behavior are strongly influenced by previous experience. However, in both groups, movement velocity of *Nlgn1*^-/-^ mice in the open-field arena was either not different (Figure 5C) or greater than WT controls (Supplemental Figure 12). This was further supported by no differences in the latencies to fall off an accelerating rotarod across repeated trials (Supplemental Figure 13) highlighting loss of Nlgn1 does not impair intrinsic motor function, therefore the observed changes in motivational measures are not due to locomotor capacity.

**Figure 5:**
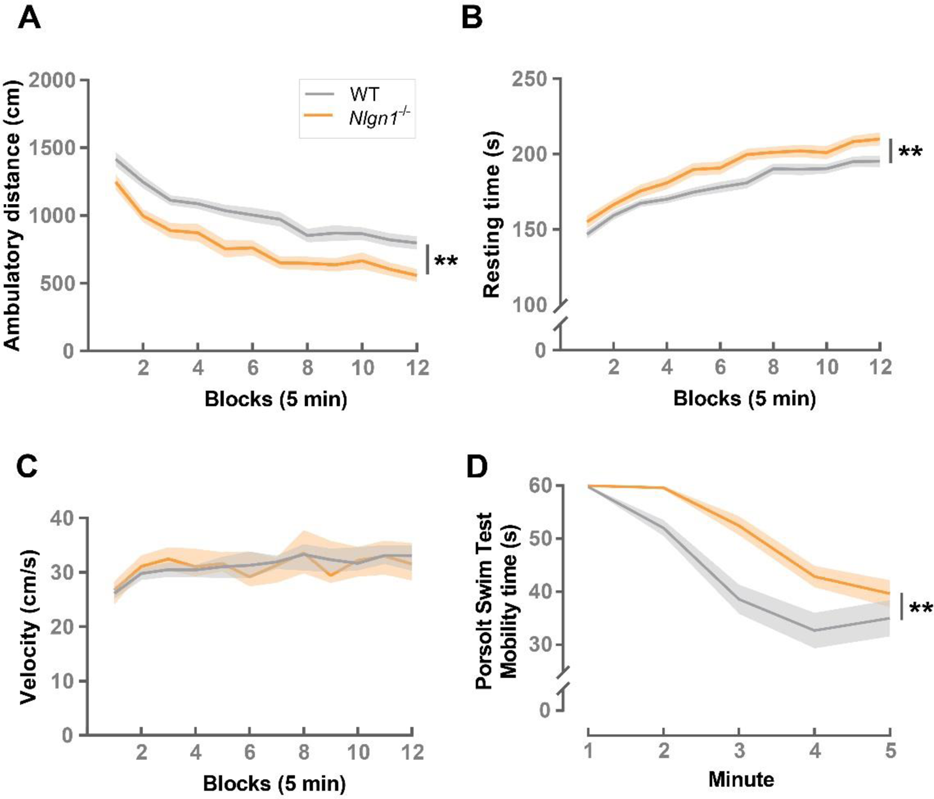
Measuring motivational behavior outside an operant environment. *Nlgn1*^-/-^ mice (with previous operant experience) show *decreased* exploration and spontaneous locomotor activity in a novel, open-field environment. **(A)** ambulatory distance (centimeters) and **(B)** resting time (seconds) (generalised linear model \*\**p* < 0.005) but no changes in **(C)** velocity (centimeters/second)(linear regression). Values represent means ± SEM. **(D)** *Nlgn1*^-/-^ mice showed *increased* mobility time (seconds) in the Porsolt Swim Test; two-way ANOVA, time bins collapsed for analysis, main effect of genotype \*\**p* < 0.005, values represent means ± SEM.

Motivation often refers to the ability to overcome physical effort to achieve a desirable outcome (43–46), but what if the desired outcome was to avoid punishment? We were therefore interested to know whether an increased aversion to physical effort to earn rewards would manifest as a decreased willingness to avoid punishment. To examine this, we next employed the Porsolt forced swim test in which the choice to swim or ‘struggle’ to escape the aversive situation of being immersed in water compared to immobility is taken to model behavioral despair (47–49). Here, we surprisingly observed that *Nlgn1*^-/-^ mice spent significantly *more* time mobile than WT controls (Figure 5D). These data suggest that the behavioral phenotype of *Nlgn1*^-/-^ mice cannot be described simply as a general reduction in the willingness to overcome effort cost.

### Convergence on a model of increased weighting on negative utilities

Theoretical accounts for the potential cognitive mechanisms underlying decreased motivation have been reported (41, 50, 51). We therefore considered the possibility that the seemingly opposing motivational phenotypes observed in reward- and punishment-driven contexts many converge on the same underlying mechanism. On reflection, in the Porsolt swim test both the fear of drowning and the effort of swimming incurs negative utilities, Nlgn1 may not only be involved in estimating the cost of physical effort, but rather more broadly important for regulating the sensitivity to punishment or negative utilities. To explore this, we sought a theoretical model to capture the key behavioral observations of *Nlgn1*^-/-^ mice across tasks including: 1) normal binary effort-matched choices between a correct and incorrect response in learning tasks (e.g., visual discrimination, paired associates, reversal); 2) fewer responses emitted on a sequential fixed-ratio task (e.g., FR5-40); and 3) higher mobility when immersed in water in the Porsolt swim test. We visualized the behavioral predictions of the theory by simulating the behavior of a simple reinforcement learning model using the three tasks mentioned, allowing us to describe precisely and unambiguously the assumptions, architecture and predictions of the theory. We compared the simulated effect of changing the parameters in the theoretical model to the effect of Nlgn1 deletion in the experimental data.

In the model, the simulated agent selects actions by comparing the net utility of each available action, sum of positive and negative utilities weighted by two separate parameters (inspired by Collins and Frank (42), Figure 6A). Importantly, these parameters affect only the weightings on learned action utilities but not the process of learning itself, allowing a potential dissociation between learning and action selection. We first considered how reducing the weighting on positive utilities (β_P_) would impact the calculation of net utilities (Supplemental Figure 14). Reducing the weighting on positive utilities promotes low-effort-low-reward actions (e.g. resting) leading to a reduction in responses made in the fixed-ratio task, consistent with our observed experimental data in *Nlgn1*^-/-^ mice. On the contrary, reducing the weighting on positive utilities renders the choice between correct and incorrect responding more random in the simulated binary choice task, because performance accuracy depends on the difference between the positive utilities associated with the correct and incorrect response. Here, we see this model does not capture the *Nlgn1*^-/-^ phenotype in the pairwise visual discrimination, object-location paired associates learning and reversal learning tasks.

Next, we considered how increasing the weighting on negative utilities (β_N_) impacts calculation of net utilities (Figure 6B). We found increasing β_N_ i) did not alter the learning curves for binary choices because correct and incorrect responding required the same physical effort; ii) reduced the number of responses in the simulated fixed-ratio tasks because the potential rewards of responding were more heavily discounted by the physical effort incurred; and iii) increased mobility in the Porsolt swim test because increasing β_N_ had a greater effect on greater punishment, under the assumption that immersion in water has a greater negative utility than the physical effort of swimming. Together, an increased weighting on negative utilities can, at least in principle, capture the divergent phenotypes across reward- and punishment-driven tasks to suggest loss of Nlgn1 may be important for balancing the weighting on positive and negative utilities in reward-cost trade-off.

**Figure 6:**
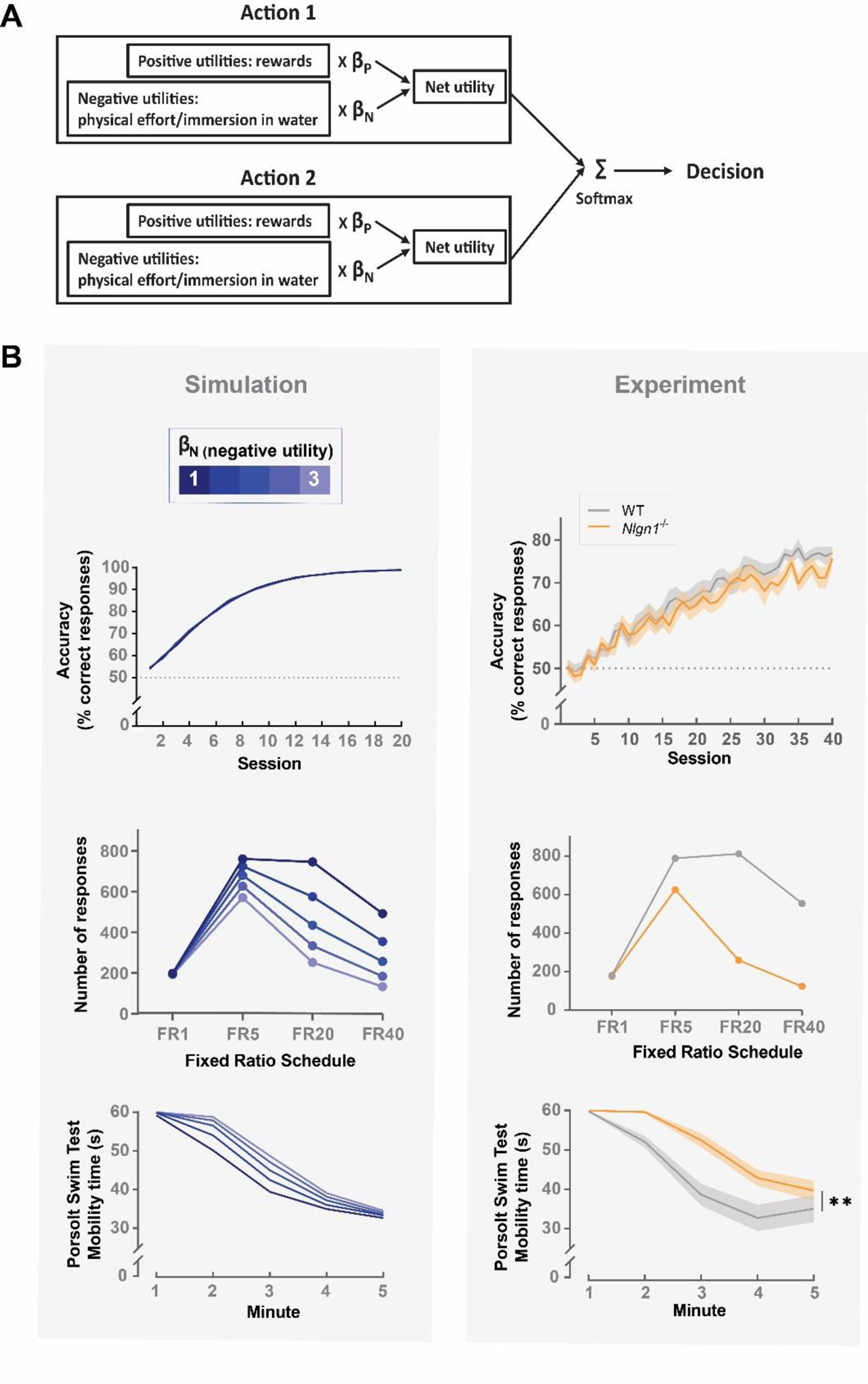
A model of increased weighting on negative utilities captures the observed *Nlgn1*^-**/-**^ **behavioral phenotype across tasks. (A)** An agent selects actions by comparing the net utility of each available action, sum of positive and negative utilities weighted by two separate parameters (β_P_ and β_N_). The model assumes different types of positive and negative utilities are under the general control of β_P_ and β_N_ (e.g., β_N_ affects both the weighting on physical effort as well as immersion in water where appropriate). Net utilities of potential actions are then fed into a softmax function for probabilistic action selection where actions with higher net utilities are selected with higher probabilities. **(B, left panel)** Increasing β_N_ has i) no impact on the choice between correct and incorrect responding in the simulated binary choice task, ii) reduces the number of responses made in the simulated fixed ratio task where the choice is between responding (high-effort action) and resting (low-effort action), and iii) increases mobility in the simulated Porsolt Swim Test where the choice is between swimming (high-effort action) and resting (low-effort action). **(B, right panel)** For comparison, our experimental data from i) Object-Location Paired Associates Learning (PAL) task, ii) Fixed Ratio task and iii) Porsolt Swim Test.

## DISCUSSION

Building on the extensive work on the molecular and signaling functions of Nlgn1 in the brain, we investigated how the loss of Nlgn1 might impact components of decision-making. We found that Nlgn1 was not required for learning complex associative structures, or the subsequent updating of learned associations. However, *Nlgn1*^-/-^ mice were consistently *less* motivated to overcome the effort cost to earn rewards across multiple task contexts that involved instrumental responding for an outcome in a free-operant environment; but *more* motivated to exert effort to avoid an inescapable aversive situation. We suggest these divergent phenotypes converge on a model of increased weighting on negative utilities, highlighting a novel valence-dependent role of Nlgn1 in reward-cost trade-off.

It is widely held that NMDAR function and long-lasting forms of synaptic plasticity are required for various forms of learning (e.g., (25-27, 29, 33, 52). Based on previous findings that modulating Nlgn1 expression robustly impairs synaptic plasticity in various brain regions and spatial memory, we initially speculated that it would likely impair the ability to learn complex associations in our various touchscreen-based tasks. Indeed, disrupting NMDAR signaling and plasticity has been shown to impair performance in these touchscreen-based tasks employing training parameters similar to our study (33, 52–55). It is therefore striking that *Nlgn1*^-/-^ mice showed normal learning and were able to acquire complex associative structures and modify learned associations. It is worth noting, however, that functional studies on synaptic proteins (including Nlgn1) have predominantly been performed *in vitro* and often limited to isolated brain regions (i.e., hippocampus) and cell-types (i.e., pyramidal neurons). The synaptic mechanisms required for the emergence of large-scale neural representations during complex learning and how these might be regulated by molecular components that structurally organize synapses therefore remains largely unknown. Technological advances in recent years allows *in vivo* monitoring of neural dynamics of large populations of cells across multiple brain regions in awake-behaving animals with high cellular specificity. These emerging approaches provide impressive depth and breadth, and offer new opportunities for probing how synaptic mechanisms influence large-scale neural dynamics underlying complex behavior.

Despite an intact capacity to acquire and flexibly update learned information, we found that loss of Nlgn1 alters motivational processing, impacting reward-cost trade-off. Action selection generally requires weighing up both the expected positive and negative consequences of available actions. We see a *reduced* willingness to overcome response effort for reward in touchscreen-based tasks. Theoretical models have been proposed to address the dissociation between learning and motivation for rewards, potentially explaining our dissociation (41, 51).

According to these theories, an animal could be less motivated to overcome the effort cost of responding yet maintain the normal ability to choose between correct and incorrect responses due to either an underestimation of the average reward rate of the environment independent of any particular action, or an overestimation of the effort costs of potential actions. However, we additionally see an *increased* willingness to exert effort to escape an aversive situation in the Porsolt swim test. One interpretation may be that this is due to an exaggerated anxiety or fear response, but we deem this to be unlikely. Previous work has shown *Nlgn1*^-/-^ mice demonstrate intact contextual and cued fear conditioning, and normal behavior on various anxiety-related assays but knockdown of Nlgn1 in the amygdala of rats impairs fear memory. Further, it is unclear to what degree the mechanisms underlying freezing behavior in Pavlovian fear conditioning overlap with immobility behavior in the Porsolt swim test.

The divergent phenotypes observed in the reward-associated touchscreen tasks and the punishment-associated Porsolt forced swim test could either be due to changes in distinct or shared underlying mechanisms involved in the decision-making processes. Our behavioral simulation data show that the seemingly opposing phenotypes could, in principle converge on an increased weighting on negative utilities. Two key questions arise from this interpretation. First, is it plausible to assume a common currency for effort cost and other forms of punishment? Conceptually, this seems reasonable since physical cost, like other forms of punishment, is a decision variable which animals should seek to minimize all else being equal (56). Empirically, functional magnetic resonance imaging (fMRI) data suggest that hemodynamic responses in select brain regions such as the anterior insula cortex correlate with both physical cost and other forms of punishment (56–60). Second, is it plausible that the brain affords dissociable neural computation and hardware for reward and punishment? It is generally accepted that there is at least partial dissociation between the neural implementation of reward and punishment (61–63). Notwithstanding the lack of a consensus, reward and punishment processing has been shown to be differentially implemented by activity (64), and identity of distinct subsets of dopaminergic neurons (65–67) or opponent dopaminergic and serotoninergic signaling (51, 68).

In summary, our findings identify Nlgn1 is essential for regulating distinct cognitive processes underlying decision-making, providing evidence of a new model for dissociating the computations underlying learning and motivational processing. These data, cultivated from our approach to comprehensively examine a battery of tests measured within the same testing environment in a controlled and comparable manner, highlight the value of a deep behavioral dissection. Human mutations in *NLGN* genes, including *NLGN1* have been reported in neurodevelopmental disorders including autism spectrum disorder (69). It is noteworthy that depression and affective symptoms are highly co-morbid with neurodevelopmental disorders, supporting the need to dissect distinct components of cognitive behavior and gain a deeper understanding of the transdiagnostic psychological processes that can be effectively used as behavioral markers of disease.

## AUTHORS’ CONTRIBUTIONS

J.L. and J.N. conceived and designed the experiments. J.L performed all experiments and analyses with assistance from J.T. in testing spontaneous locomotor activity and Porsolt swim test. J.L. and J.N. wrote the manuscript.

## ACKNOWLEDGEMENTS

This work was supported by an Australian Research Council Future Fellowship (FT140101327) and Brain and Behavioral Foundation (NARSAD) Young Investigator Award to J.N., Australian Postgraduate Research Training Award to J.L, and a National Health and Medical Research Council Project Grant (APP1083334).

We thank Prof. Nils Brose (Max Planck Institute for Experimental Medicine, Göttingen Germany) for providing breeding founders of *Nlgn1^+/-^* mice, Prof. Leonid Churilov for statistical advice, Shana Schokman and Clara Lee for genotyping, Brittany Cuic, Leah Payne and Florey Core Animal Services for mouse husbandry, and Debbie Maizels for manuscript artwork contribution.

## METHODS

### Animals and Housing

*Nlgn1*^-/-^ mice were obtained from Prof. Nils Brose, generated by homologous recombination of embryonic stem cells deleting exon sequences covering the translational start site and 546 bp of 5’ coding sequence of the murine *Nlgn1* gene (70), and backcrossed more than 10 generations on a C57BL/6 background. *Nlgn1*^-/-^ mice and WT littermate matched controls were generated by mating heterozygous females and males. Mice were weaned at 3-4 weeks of age and housed in mixed genotype groups of 2-4 per cage with food and water available *ad libitum*. Bedding consisted of sawdust chips 2cm deep and tissue paper for nesting material. At ∼10 weeks of age, mice were moved from individually ventilated cages to open-top cages in a humidity and temperature-controlled holding room maintained on a 12:12-hour reversed light/dark cycle (lights off at 07:00). Mice were acclimatized to these conditions for a minimum of one week prior to handing. Pre-training began at ∼12 weeks of age. All behavioral testing was conducted during the dark active phase of the cycle, with the experimenter blinded to genotype during behavioral testing. All procedures were approved by the Florey Institute of Neuroscience and Mental Health Animal Ethics Committee.

### Cohorts of mice used for behavioral testing

A total of 6 cohorts of mice were used in the present study. <u>Cohort 1</u> (WT: n=12 female/n=15 male; *Nlgn1*^-/-^: n=13 female/n=13 male) were tested in the pairwise visual discrimination, reversal learning, object-location paired associates learning and extinction learning tasks. When a single cohort of animals were tested on multiple touchscreen-based tasks, mice were placed back on free-feeding for ∼2 weeks and weights updated prior to food-restriction and commencing next task. <u>Cohort 2</u> (WT: n=14 female/n=14 male; *Nlgn1*^-/-^: n=14 female/n=17 male) were tested in the fixed ratio task (FR1-40) with strawberry milk rewards and fixed ratio 20 (FR20) task with water rewards. <u>Cohort 3</u> (WT: n=11 female/n=12 male; *Nlgn1*^-/-^: n=7 female/n=12 male) and <u>Cohort 4</u> (WT: n=6 female/n=4 male; *Nlgn1*^-/-^: n=6 female/n=4 male) were tested in the progressive ratio task, spontaneous locomotor activity and accelerating rotarod tests. <u>Cohort 5</u> (WT: n=13 female/n=10 male; *Nlgn1*^-/-^: n=10 female/n=12 male) were tested in the Porsolt forced swim test following ∼2 weeks simple operant training for a different study. <u>Cohort 6</u> (WT: n=13 female/n=16male; *Nlgn1*^-/-^: n=11 female/n=16 male) were experimentally naive and tested for spontaneous locomotor activity. For all non-touchscreen-based tasks, mice were not food restricted when tested.

### Rodent Touchscreen Operant Tasks

#### Apparatus

Touchscreen testing was conducted in the Bussey-Saksida mouse touchscreen operant system (Campden Instruments Ltd, UK). Stimulus presentation, task parameters and data recording were controlled through Whisker Server and ABET II Touch software (Campden Instruments Ltd, UK).

#### Touchscreen pre-training

Pre-training and food restriction were conducted as previously described (34, 35, 39). Before testing, mice were first food restricted to 85 – 90% free-feeding body weight. Mice were then trained through five phases or instrumental conditioning to learn to selectively nose-poke stimuli displayed on the touchscreen in order to obtain a liquid reward (strawberry milk, Devondale, Australia). Mice were required to reach a set performance criterion for each phase before advancing to the next phase. Briefly, mice were habituated (phase 1, Habituation) to the touchscreen chamber and to consuming liquid rewards from the reward magazine or receptacle for two 30-minute sessions (criterion = consume 200µl of liquid reward freely available in the reward receptacle at each session). For phases 2–5, a trial did not advance until the reward was consumed. In phase 2 (Initial Touch) or the Pavlovian stage, a single visual stimulus was displayed on the screen for 30 s, after which the disappearance of the stimulus coincided with delivery of a reward (20 μl), presentation of a tone and illumination of the reward receptacle (criterion = 30 trials in 60 min). A nose-poke response to the stimulus during the 30s window was rewarded with 3 times the reward amount to encourage responding. In phase 3 (Must Touch), mice had to nose-poke visual stimuli displayed on the screen to obtain a reward (criterion = 30 trials in 60 min). Mice then learned to initiate a new trial with a head entry into the reward receptacle (phase 4, Must Initiate, criterion = 30 trials in 60 min). In phase 5, responses at a blank part of the screen during stimulus presentation produced a 5-s timeout (signaled by illumination of the house light and no delivery of reward) to discourage indiscriminate responding (criterion = 21/30 correct responses in 60 min on 2 consecutive days). If another response to a blank part of the screen during stimulus presentation was made, there was a 5s inter-trial interval (ITI), and then the same trial was repeated (the same stimulus presented in the same screen location, termed a correction trial) until the mouse made a correct response. Therefore Phases 2-5 consisted of 30 trials (pseudorandom first-presentation), and Phase 5 also included an unlimited number of correction trials.

#### Pairwise visual discrimination and reversal learning

The pairwise visual discrimination (PD) and reversal learning (RL) tasks were conducted like that previously described (34, 35, 39). Briefly, mice were trained to discriminate between two novel, equiluminescent visual stimuli (left and right diagonal stripes) displayed pseudorandomly across two locations with equal number of appearances at each location. Response to one stimulus resulted in reward delivery (S+, correct response), followed by a pseudorandom trial (maximum 30 per session); response to the other stimulus resulted in a 5-second timeout, illumination of the house light followed by a correction trial. The same stimulus configuration was presented on correction trials until a correct response was made and a reward was delivered. Correction trials were not counted towards the trial limit or percentage of correct responses of a session. The designation of S+ and S- was counterbalanced within genotype and sex groups. Mice were trained to an acquisition criterion of ≥80% correct responses on two consecutive sessions. Following the acquisition of the visual discrimination task, mice were immediately moved on to the reversal leaning task, where the previously acquired reward contingencies were reversed. Reversal learning was assessed across 20 sessions.

#### Object location paired associates learning

The object-location paired associates learning (PAL) task was conducted as previously described (34, 35). Briefly, mice were trained to acquire reward associations jointly defined by visual stimuli (flower, plane and spider) and their assigned correct spatial locations on the touchscreen (left, center and right, respectively). For each trial, only two objects were presented: one object in its correct location (S+) and the other object in one of two incorrect locations (S−); therefore, there were six possible trial types. A nose-poke to the S+ resulted in delivery of a reward followed by a pseudorandom trial (maximum 36 per session), and incorrect responses resulted in a 5 s time-out followed by correction trial. Visuospatial learning in the PAL task was assessed across 40 sessions.

#### Instrumental extinction learning

The instrumental extinction learning task was conducted similar to that previously described (35, 39). Mice were first trained to make a nose-poke response to a single white square displayed on the touchscreen for a reward until reaching a set acquisition criterion (30 trials in <12.5 min on five consecutive sessions). Following acquisition, instrumental extinction was assessed where responses were no longer rewarded (30 trials per session tested across 6 sessions). During extinction, the visual stimulus was displayed for 10 s on each trial and animals could either make a response or an omission.

#### Progressive ratio

Details on testing the touchscreen-based progressive ratio task has been described previously (71). Briefly, naive mice first underwent phases 1 and 2 of touchscreen pre-training, followed by one session each of fixed ratio schedules of 1 (FR1), FR2, FR3 and three sessions of FR5 training where a fixed number of nose-pokes (1, 2, 3 and 5 respectively) were required for a reward. Mice were required to complete 30 trials in 60 minutes in each of the FR sessions (criterion). Once training criterion was reached, mice advanced to the progressive ratio stage where the number of nose-poke responses required to obtain a reward incremented by 4 after every trial (1, 5, 9, 13 etc.,). If no responses to the touchscreen or entries to the reward receptacle were detected for 5 min, the session ended and the animal removed from the chamber.

#### Fixed ratios

Naive mice first underwent phases 1 and 2 of touchscreen pre-training followed by three sessions of FR1 and had to complete 30 trials within a 60 min-session before advancing. During the next serial FR test stage, mice were given 60 min per session to make as many responses as they were willing to, and sessions did not terminate due to inactivity. Mice were tested on three sessions of FR1, FR5, FR20 and FR40 sequentially.

#### Fixed ratio with water rewards

Following the serial FR testing, mice were water-restricted with access to water limited to one hour per day. Water-restricted body weights were maintained between 85 – 90% of free-feeding body weight. Mice were tested on a FR20 schedule where 20 nose-poke responses were required to deliver a water reward (20µl) for three sessions. After each session, mice were returned to home cage and given one-hour free access to water.

### Non-Operant Behavioral Tests

#### Spontaneous locomotor activity

Mice were assessed for spontaneous locomotor activity in a novel open-field arena (Med Associates, St. Albans, VT, USA) using the Activity Monitor system and software (Med Associates, St. Albans, VT, USA) in a 60-minute session in darkness (to minimize stress).

#### Accelerating rotarod

For motor coordination and learning on the accelerating rotarod, mice were exposed to three 5-min trials across 3 consecutive days (9 trials in total). Mice were placed on a rotating rod (Ugo Basile, Gemonio, VA, Italy) that accelerated from 4 to 40 rpm and latency to fall off was manually recorded. Testing was conducted under low lighting settings (20 lux red light).

#### Porsolt forced swim test

Mice were individually placed into a beaker (13cm diameter) with 1.6L of water (23 – 25°C) for a single 5 min session. Each session was video-recorded, and mobility time was scored using the ForcedSwimScan software (CleverSys Inc, VA, USA).

### Data Analysis

Effect size of task variables (genotype, sex, session, stimulus location etc.,) on performance measures (accuracy, latencies etc.,) were estimated together with 95% confidence intervals (CI) and statistical significance using various two-level mixed-effect general or generalized linear models (StataCorp, TX, USA). Mice was treated as level-2 clusters and random intercepts. Binary performance measures (correct/incorrect response, response/omission) were analyzed trial-by-trial using the GLLAMM (generalized linear latent and mixed models) program (36) with a log link, whereby the effects of task variables were expressed as odds ratios with an odds ratio of 1 indicating no effect (e.g., an effect of session > 1 indicates response accuracy improves over sessions). Latency data were analyzed using quantile regressions with robust and clustered standard errors (72) from the 0.05- to 0.95-quantile at 0.05 steps to allow distribution-wide comparisons, whereby effects of task variables were expressed as latency difference with 0 indicating no effect (e.g., an effect of genotype > 0 for a given quantile indicates *Nlgn1*^-/-^ mice have longer latencies).

For spontaneous locomotor activity, ambulatory distance and mobility time (300s – resting time) were analyzed with GLLAMM with a log link. Other performance measures were analyzed using the a mixed-effects linear model if the performance measures were normally distributed or median regressions otherwise (72).

To analyze the effect of correction trials and reoccurring pseudorandom trials on accuracy, two additional binary variables were included in the models indicating whether a trial is a correction trial/reoccurring trial (correction trials were excluded in estimating the effect of reoccurring pseudorandom trials). Heteroskedasticity-robust standard errors adjusted for clustering within animals were used for all analyses.

### Behavior Simulation

#### Agent

A simple reinforcement learning agent learned the utility of an action following the classic Rescorla-Wagner rule (73):

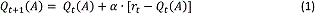

Here, *Q*_t_(*A*) is the learned utility of a given action *A* on trial *t*, *⍺* is the learning rate, *r_t_* is the reinforcement received on trial *t*. Actions can have both positive and negative utilities (e.g., responding may result in rewards but also incurs effort). The net utility of a given action is given by the linear combination of its positive and negative utilities, the relative importance of which is controlled independently by *β*_P_ and *β*_N_ respectively:

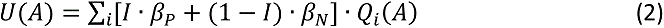

Here, *U(A)* is the net utility of action *A*, Q_i_(*A*) is the different positive or negative utilities of *A, I* is the indicator function:

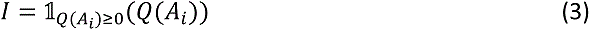

Such that,

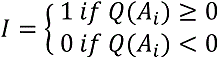

Action selection is given by a softmax function on the net utilities of potential actions

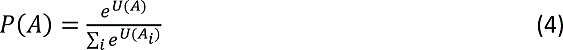

Here, P(*A*) is the probability of choosing action *A*, which depends on the net utility of *A* compared to that of alternative actions. For all our simulations, a choice is made between only two actions.

#### Simulations

##### Binary choice (two-armed bandit) task

The reinforcement learning agent learned to choose between a correct and an incorrect response for 30 trials per session across 20 sessions. The correct response was always rewarded, and the incorrect response never rewarded. Both correct and incorrect responding incurred a negative utility of −1 representing the physical effort of responding.

##### Serial fixed ratio task

The agent was trained sequentially through FR1, 5, 20 and 40 for three sessions on each ratio requirement where it chose between responding or resting. Responding resulted in a reward if the ratio requirement was met (positive utility) and incurred a negative utility of −1 representing the physical effort. Resting results in no reward but incurs a much smaller effort-related negative utility of −0.2. Note that the designation of the alternative action as resting is arbitrary. The general idea is that an animal chooses between responding and some other low-reward-low-effort actions. Time elapsed as the agent chose to either respond or rest, the session ended after 2700 timesteps roughly corresponding to a 2700-second or 45-minute session.

##### Porsolt swim test

The forced swim test was simulated as a choice between swimming and resting. Swimming was initialized with a utility of 0 representing that the agent initially believed that swimming will lead to a neutral outcome, and a utility of −1 represent the effort of swimming. Resting had a large negative utility of −10 representing the possibility of drowning but incurred no effort. Every time the agent chose to swim, it received a reinforcement of −9 thereby gradually learned by equation (1) that swimming did not markedly improve the situation therefore reduced mobility over time. For simplicity, the agent made 300 decisions over 300 timesteps roughly corresponding to a 5-min session.

Note the logic of the proposed model does not depend on the specific values of task parameters used for the simulations, which were chosen so that the behavioral simulations are quantitatively similar to the experimental data.

## Supplemental Figures

**Supplemental Figure 1:**
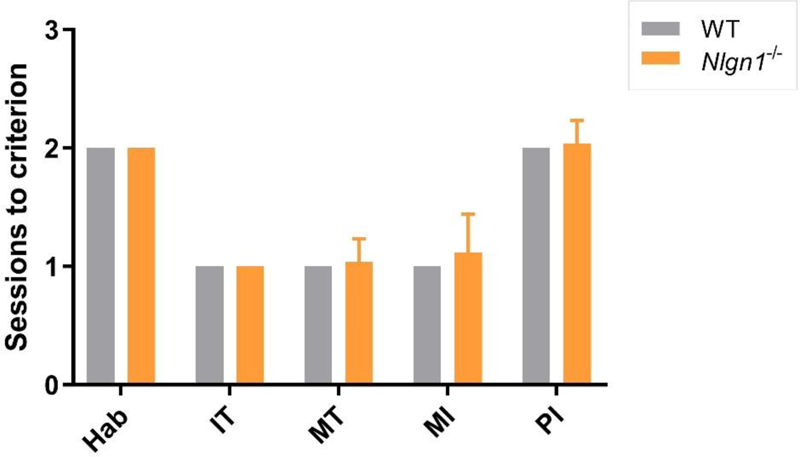
*Nlgn1*^-/-^ and WT mice required similar number of sessions to acquire touchscreen pre-training. Mice were trained through five phases: (i) Habituation, Hab; ii) Initial touch, IT; iii) Must touch, MT; iv) Must initiate, MI; v) Punish Incorrect, PI to initiate trials and selectively nose-poke visual stimuli displayed on the touchscreen in order to obtain rewards (Methods). A criterion for each phase had to be reached before advancing to the next phase.

**Supplemental Figure 2:**
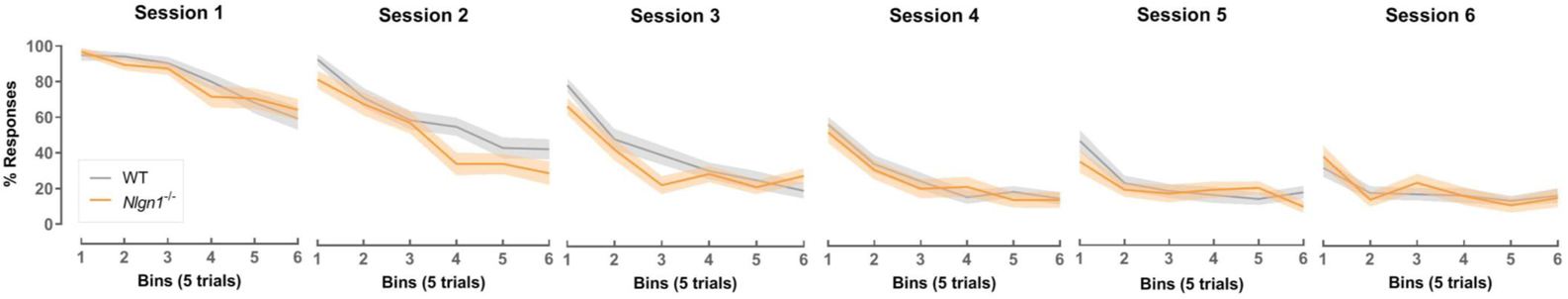
Instrumental extinction learning curves. Percentages of responses within a session (blocks of 5 trials) and across sessions in the instrumental extinction learning task. Values represent means ± SEM.

**Supplemental Figure 3:**
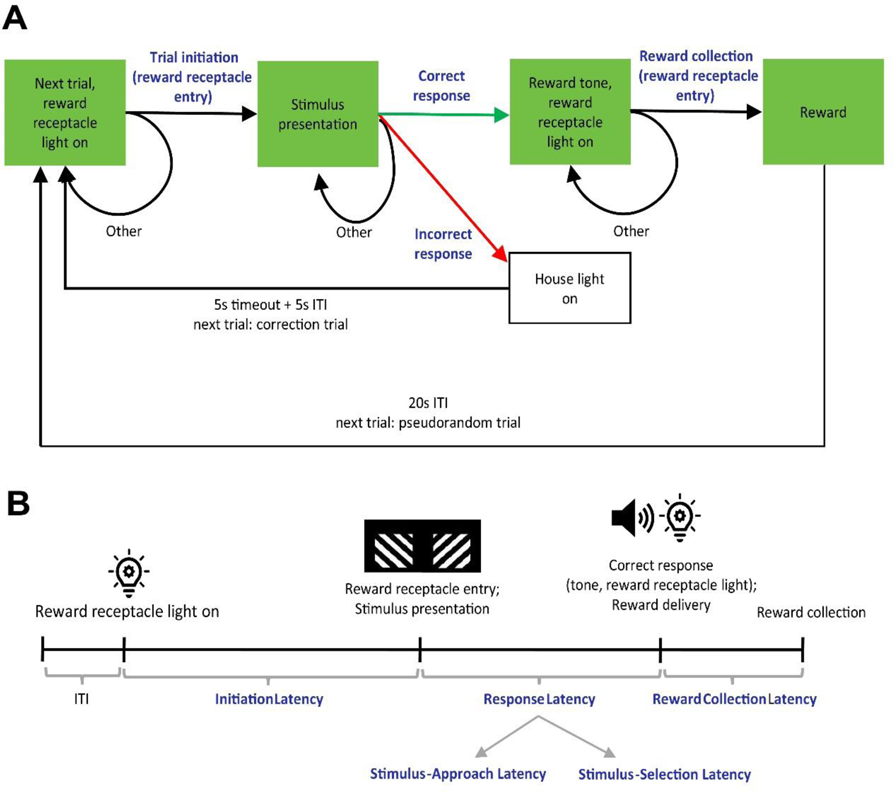
Task dynamics and latency measures of the pairwise visual discrimination (PD), reversal learning (RL) and object-location paired associates learning (PAL) tasks. (A) Task-state transitions. Green boxes represent task states and associated cues. Arrows represent transitions to the next state or staying in the current state. Blue text labels represent actions leading to transitions. ITI: inter-trial intervals. (B) Latency measurements. Illumination of the reward receptacle light signals the availability of the next trial after an ITI. Initiation latency measures the time from the end of ITI to trial initiation by head entry into the reward receptacle. Head entry triggers the presentation of stimuli. Stimulus-approach latency measures the time from exiting the magazine to arriving in front of the touchscreen. Stimulus-selection latency measures the time from arriving in front of the touchscreen to nose-poking one of the stimuli. If the response is correct, a reward tone and the reward receptacle light signal the delivery of a reward. Reward collection latency measures the time from delivery of the reward tone to head entry into the reward receptacle.

**Supplemental Figure 4:**
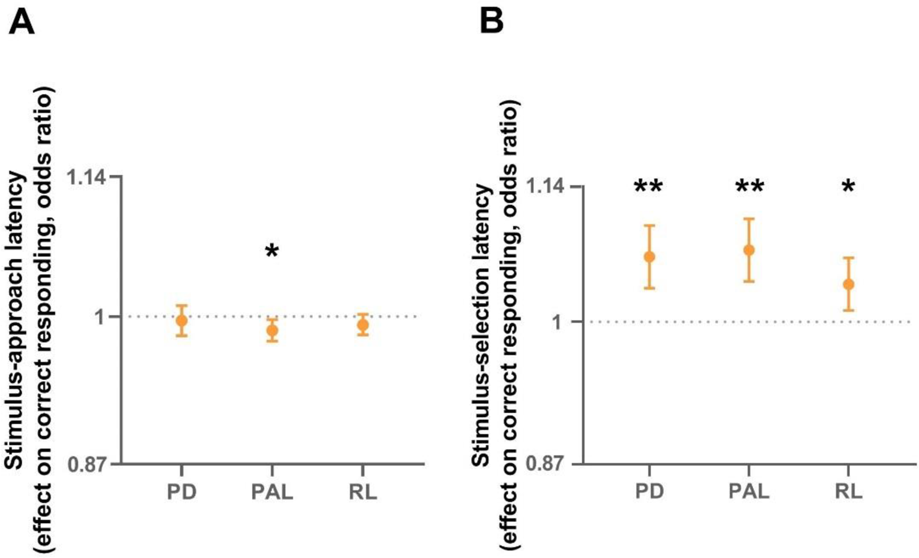
Stimulus-selection latency positively predicts response accuracy but not stimulus-approach latency. (A) Stimulus-approach latency does not positively predict response accuracy across pairwise visual discrimination (PD), object-location paired associates learning (PAL) and reversal learning (RL). However, **(B)** stimulus-selection latency positively predicted increased response accuracy across PD, PAL and RL. See Supplemental Table 1 for statistics. Logistic regression, \**p* < 0.05, \*\**p* < 0.005 significantly different from 1, values represent odds ratio ± 95% CI.

**Supplemental Figure 5:**
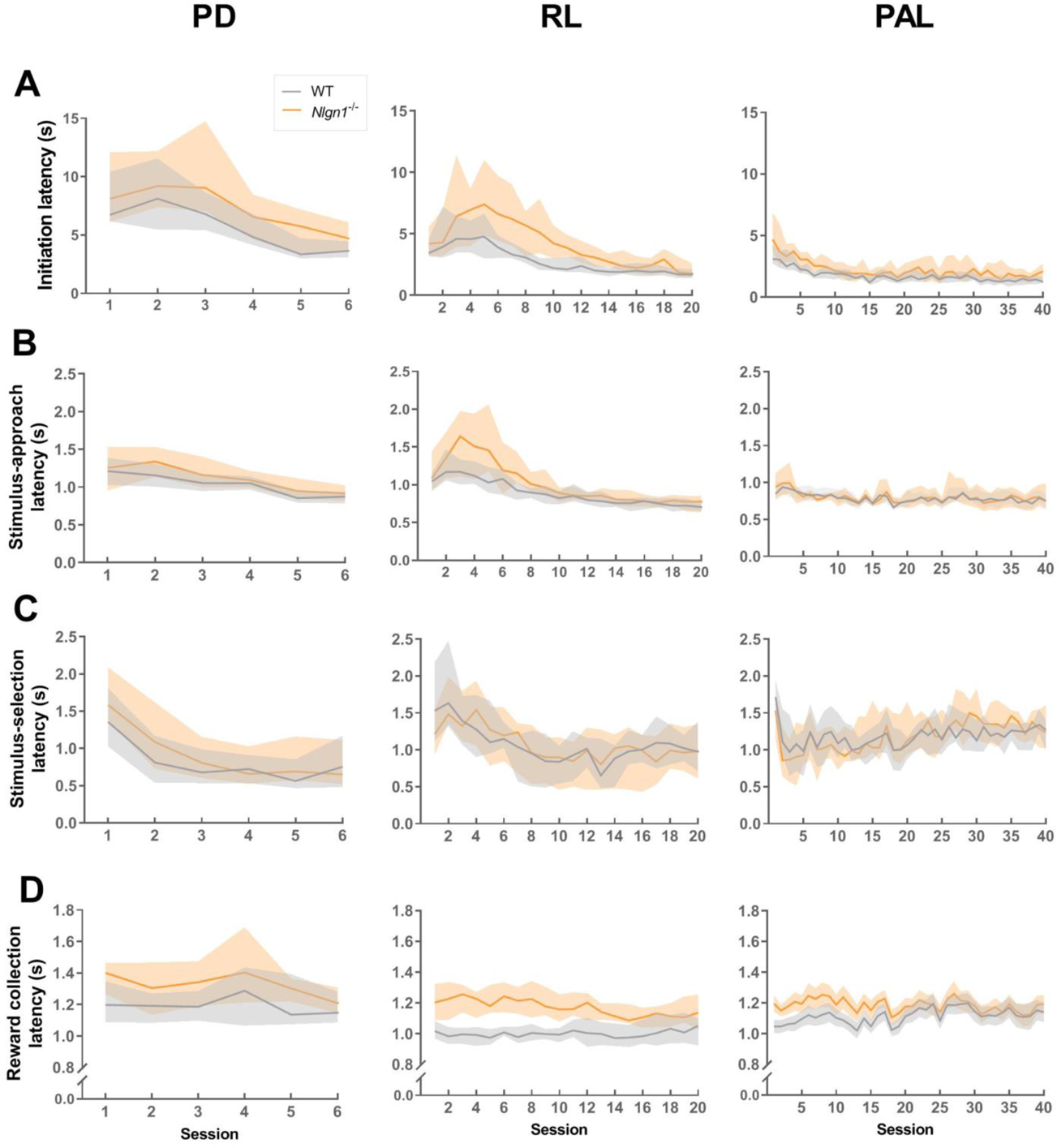
Latency learning curves. Learning curves showing the median **(A)** initiation latency, **(B)** stimulus-approach latency, **(C)** stimulus-selection latency and **(D)** reward collection latency across pairwise visual discrimination (PD), reversal learning (RL) and object-location paired associates learning (PAL) tasks. Tasks arranged in columns (left to right) in order of training. Only the first 6 sessions of PD containing all mice visualised (prior to some mice subsequently advancing after reaching criterion). Values represents median ± 95% CI.

**Supplemental Figure 6:**
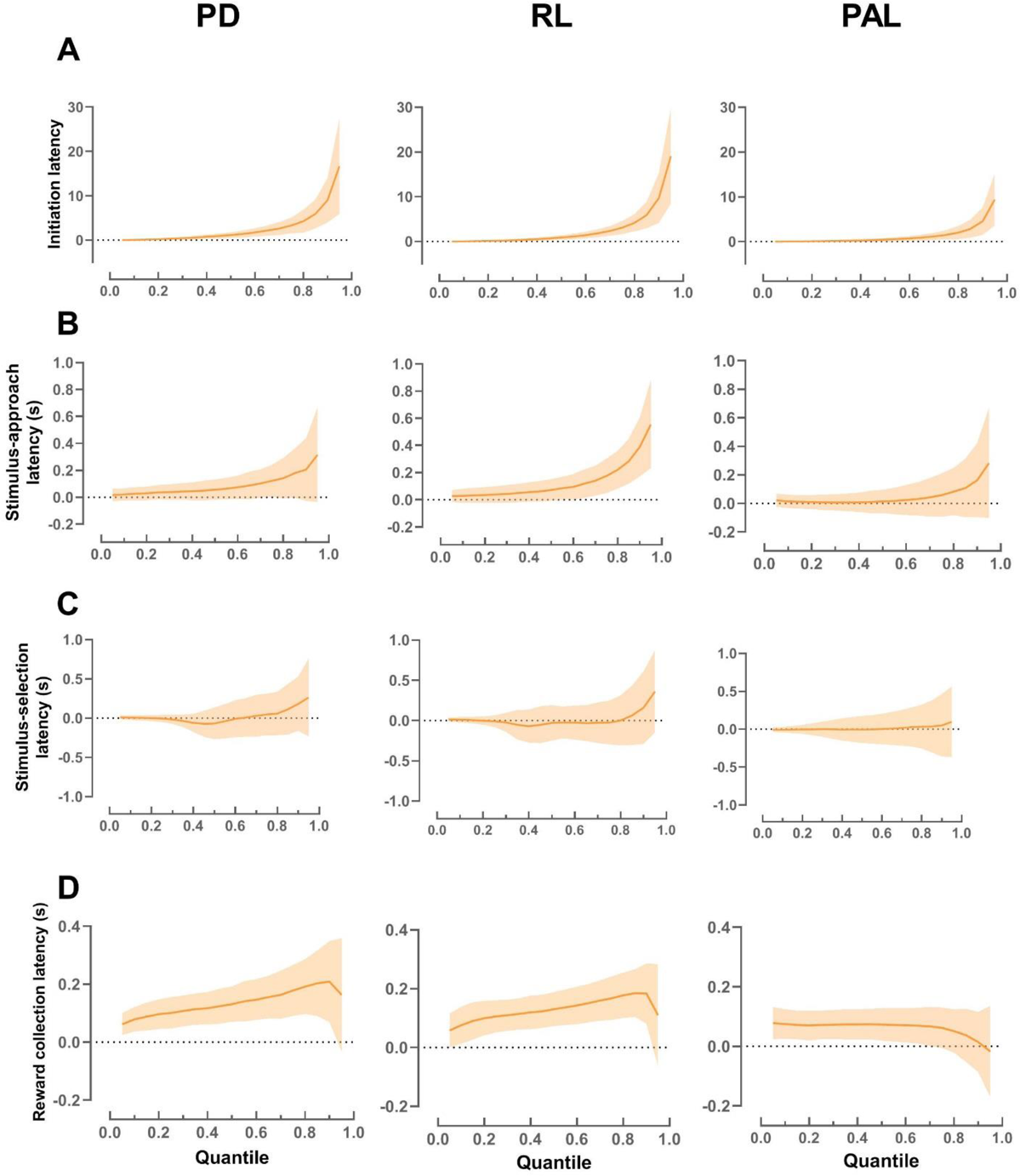
Effect of genotype on latency measures. Quantile regressions on latencies from the 0.05^th^ to the 0.95^th^ quantile at steps of 0.05. *Nlgn1*^-/-^ mice showed a general increase in **(A)** initiation latency, **(B)** stimulus-approach latency and **(D**) reward collection latency but genotype had very little effect on **(C)** stimulus-selection latency across pairwise visual discrimination (PD), reversal learning (RL) and object-location paired associates learning (PAL) tasks. Quantile regression values represent estimated latency difference between *Nlgn1*^-/-^ and WT mice ± 95% CI.

**Supplemental Figure 7:**
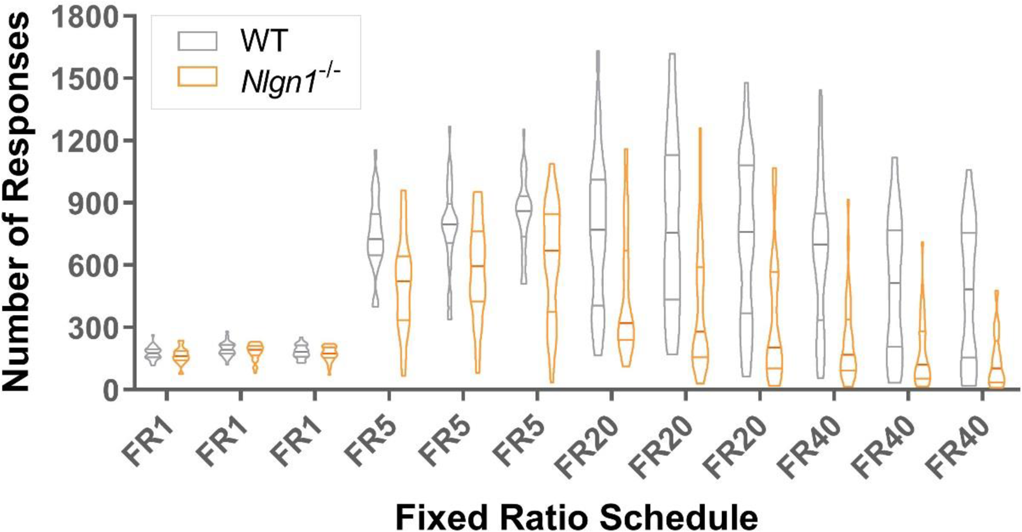
Number of responses made in Fixed Ratio (FR) task. Three sessions at each ratio requirement. Horizontal lines indicate 1^st^, 2^nd^ (median) and 3^rd^ quartiles.

**Supplemental Figure 8:**
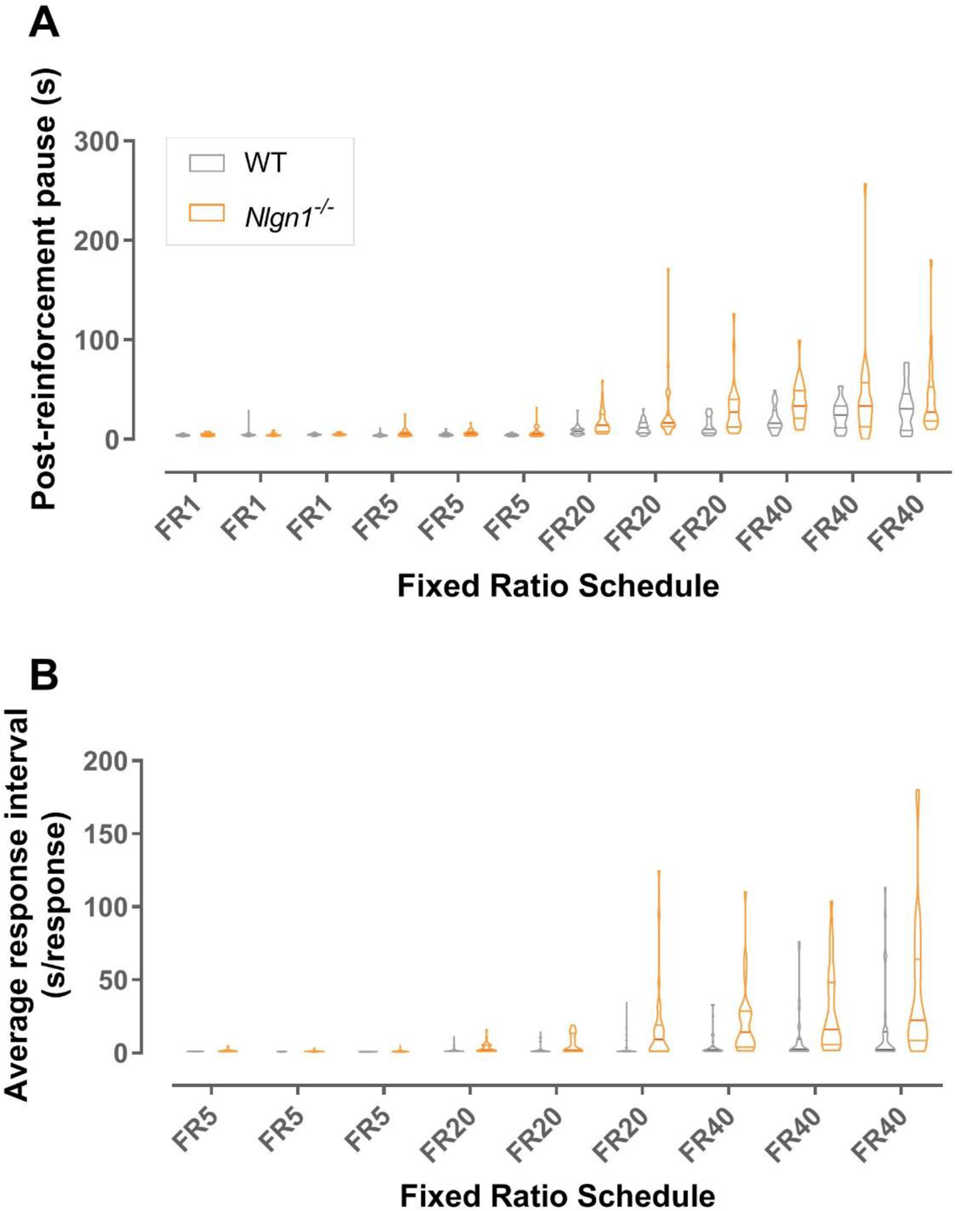
Post-reinforcement pause and average response interval in Fixed Ratio (FR) task. Three sessions at each ratio requirement. **(A)** Post-reinforcement pause: time to the first response after consuming a reward. Note data could not be gathered from animals that completed <1 trial. **(B)** Average response interval: time spent per response after the animal has made the first response of a trial. Horizontal lines indicate 1^st^, 2^nd^ (median) and 3^rd^ quartiles.

**Supplemental Figure 9:**
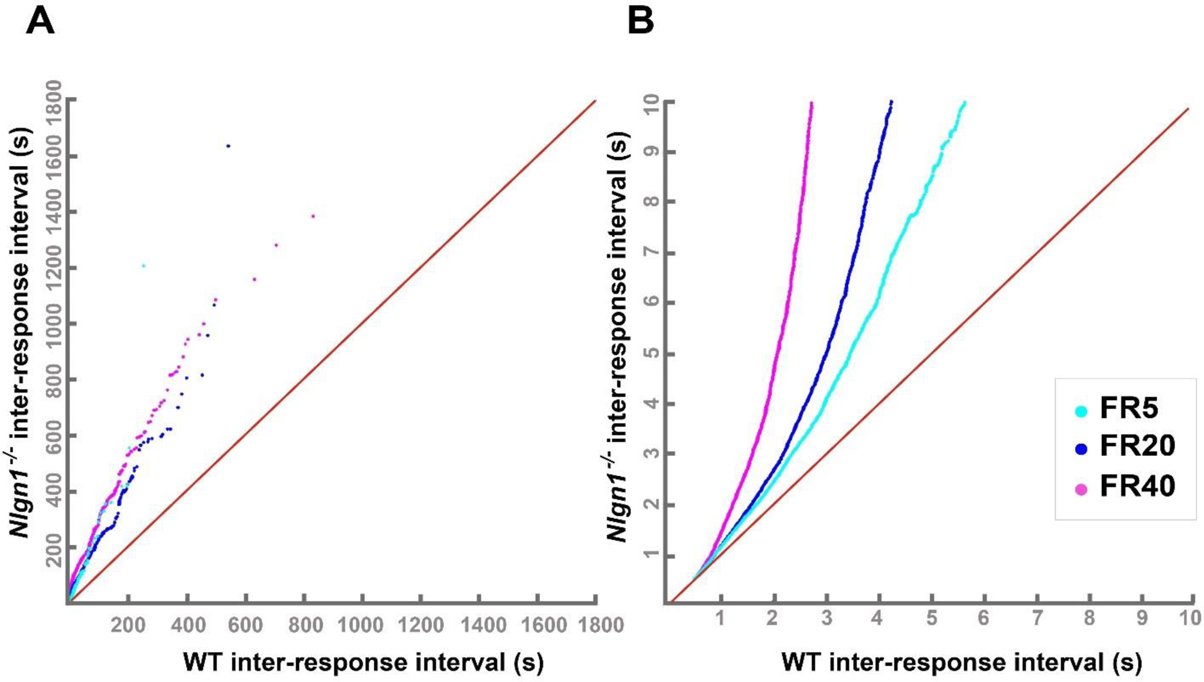
Response-by-response comparison of inter-response intervals between *Nlgn1*^-/-^ and WT mice. Quantile-quantile plots comparing the distribution of inter-response intervals (IRI). *Nlgn1*^-/-^ distribution is shifted towards longer IRI. Red line indicates identical distribution (y=x).

**Supplemental figure 10:**
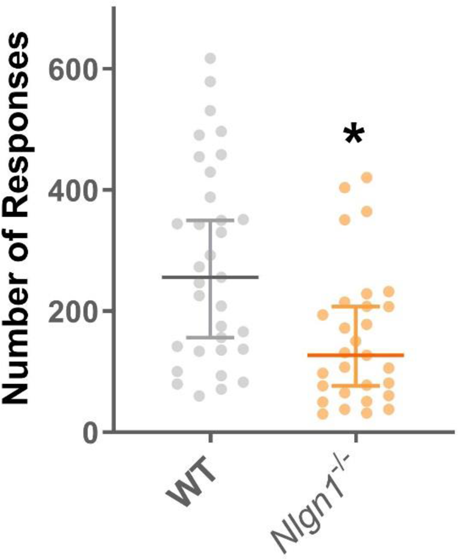
Total number of responses in progressive ratio task. *Nlgn1*^-/-^ mice (naive cohort) made fewer responses than WT controls. Quantile regression (median), \**p* < 0.05, values represent median ± 95% CI.

**Supplemental Figure 11:**
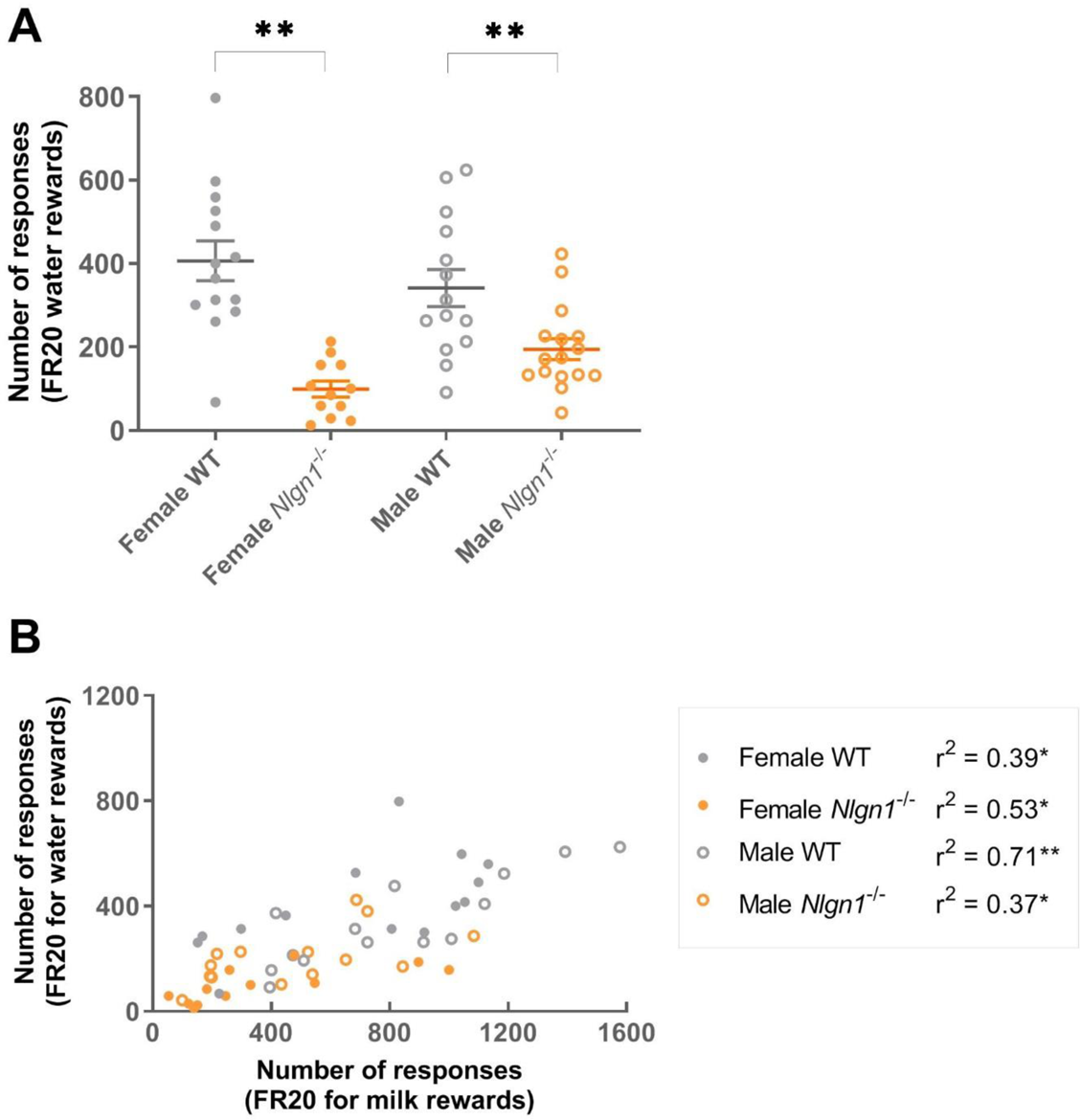
*Nlgn1*^-/-^ mice show decreased number of responses for water rewards. **(A)** Both male and female *Nlgn1*^-/-^ made significantly fewer responses than WT controls, but there was also a significant genotype x sex interaction with female *Nlgn1*^-/-^ mice making even less responses than male *Nlgn1*^-/-^ mice (see Supplemental Table 1 for statistics). **B**) Positive correlations between number of responses mice made for milk and water rewards were significant for each sex and genotype group. Linear regression, \**p* < 0.05, \*\**p* < 0.005

**Supplemental Figure 12:**
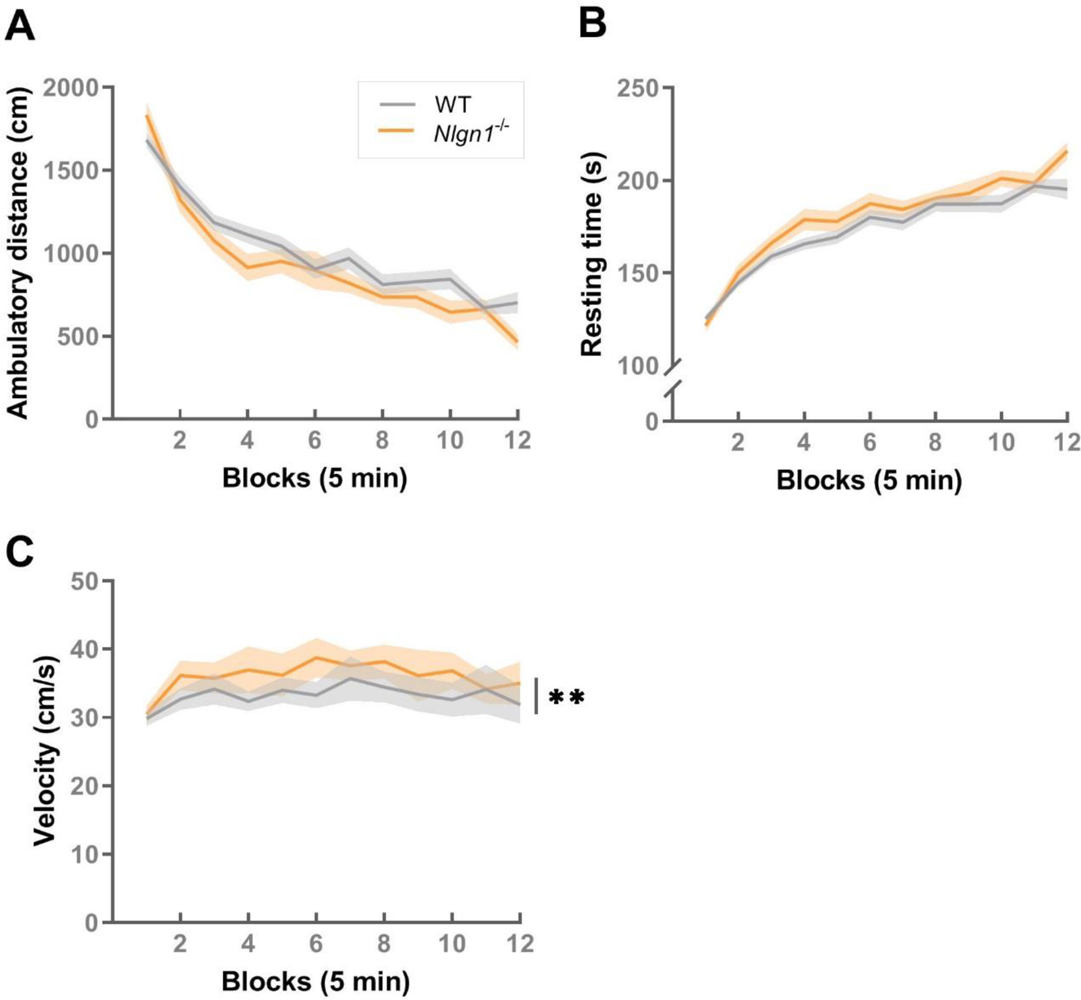
Experimentally naive *Nlgn1*^-/-^ mice show subtle changes in exploration and spontaneous locomotor activity in a novel, open-field environment. (A) Ambulatory distance (centimeters) and (B) resting time (seconds) showed no significant effect of genotype, but a significant genotype x time interaction (*p* < 0.05, generalized linear model). (C) *Nlgn1*^-/-^ mice also showed higher ambulatory velocity (centimeters/second) (effect of genotype \*\**p* < 0.005, linear regression). Values represent means ± SEM.

**Supplemental Figure 13:**
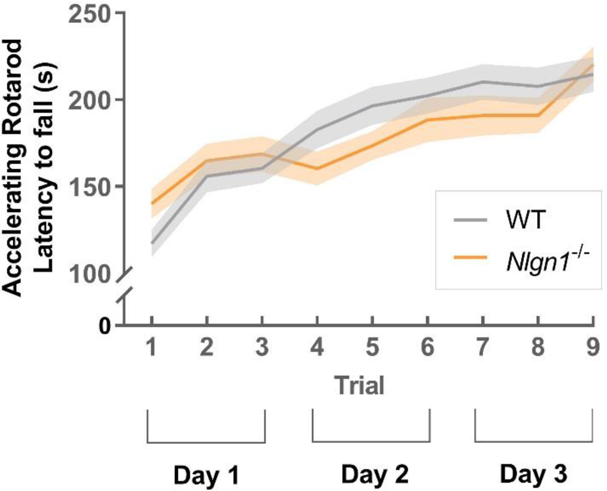
***Nlgn1*^-/-^ mice displayed normal motor coordination and learning on the accelerating rotarod test.** No differences between genotypes in the latency to fall off the accelerating rotarod on a series of three 5-min trials tested across three consecutive days (9 trials in total). Linear regression, values represent means ± SEM.

**Supplemental Figure 14:**
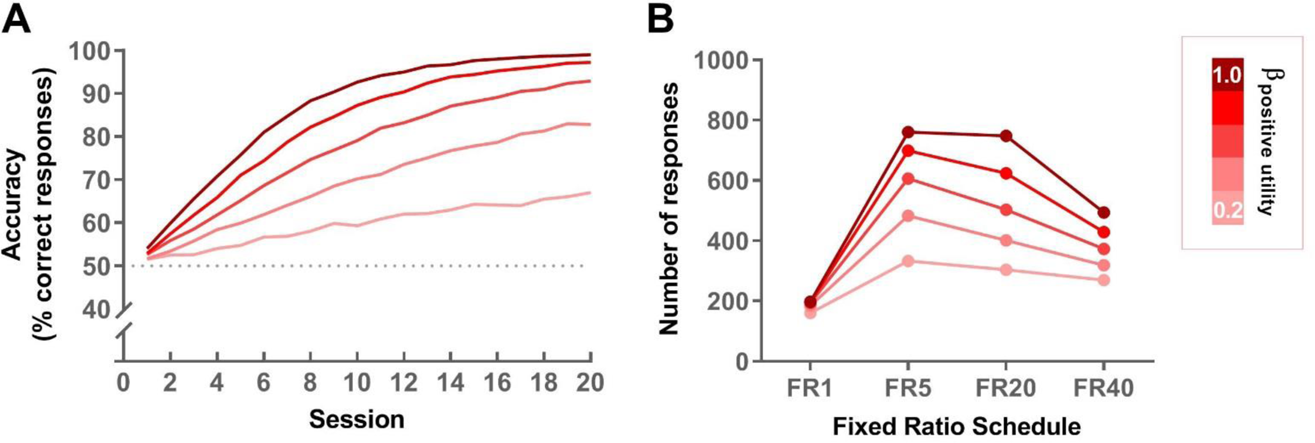
**Simulated effect of decreasing the weighting on positive utilities (β**_P_**) in the calculation of net utilities. A**) Decreasing β_P_ increases randomness in the choice between correct and incorrect responding in the simulated binary choice task leading to a flatter learning curve. **B**) Decreasing β_p_ reduces number of responses in the simulated fixed ratio task.

---

**Supplemental Table 1:**
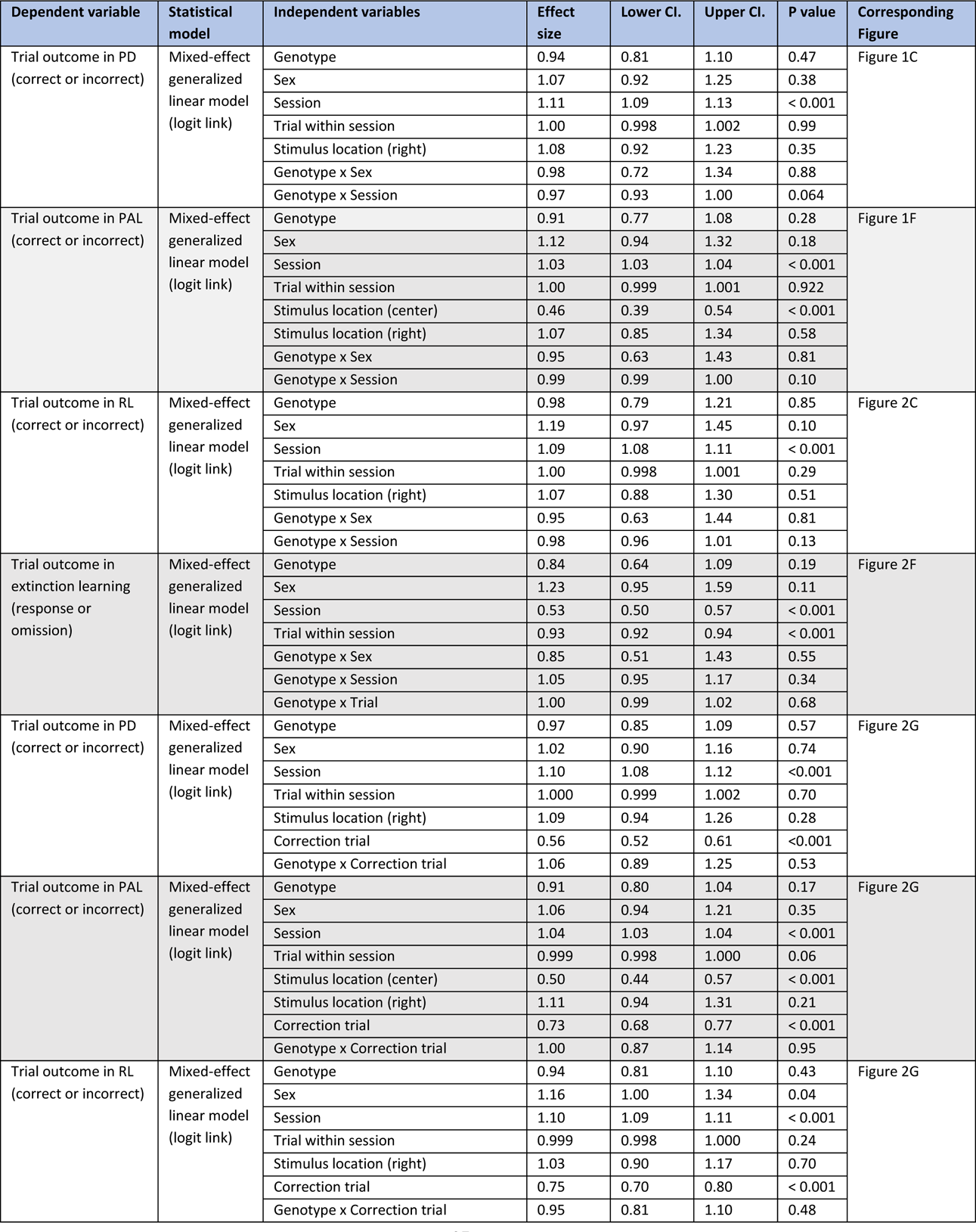
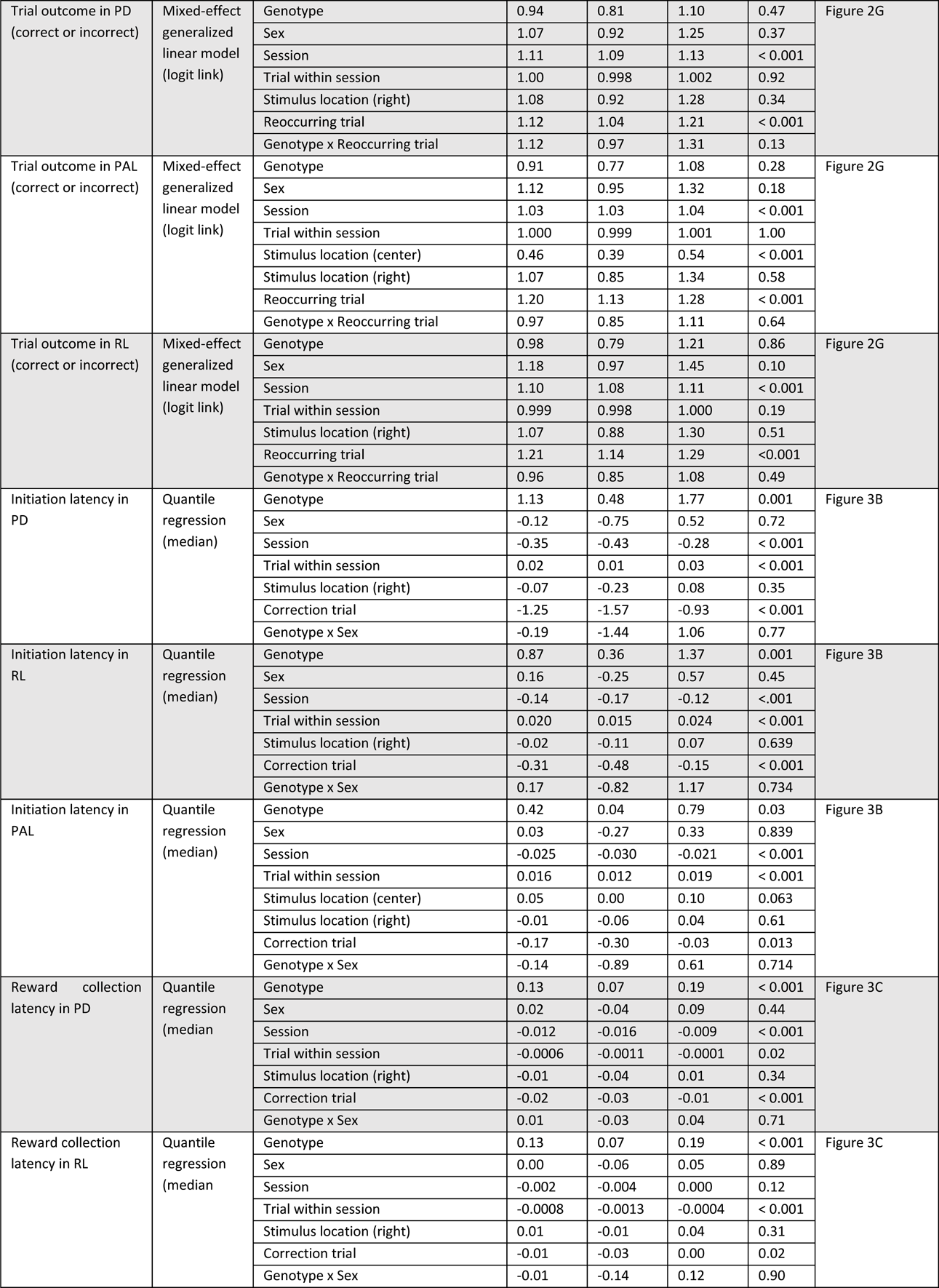
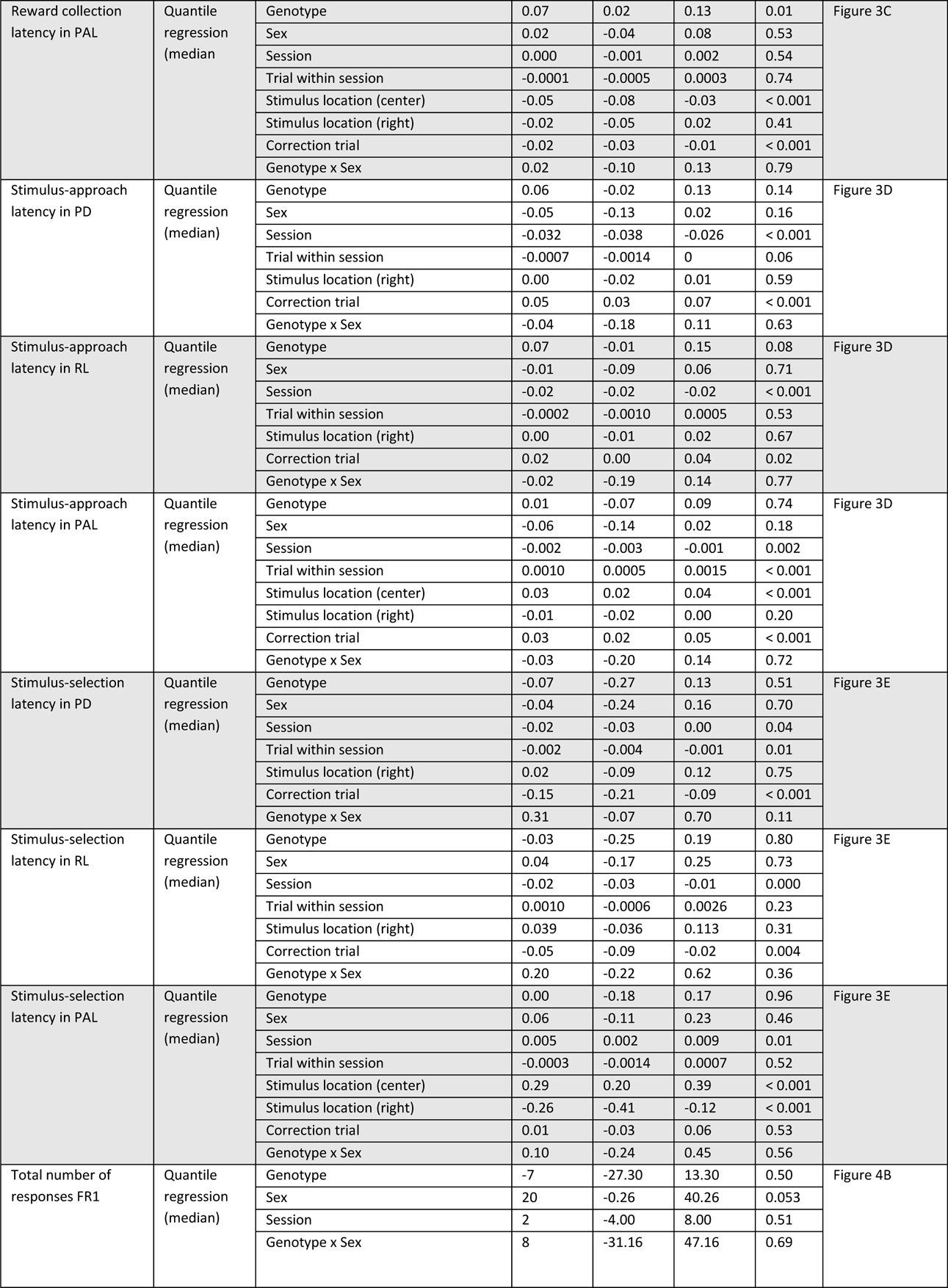
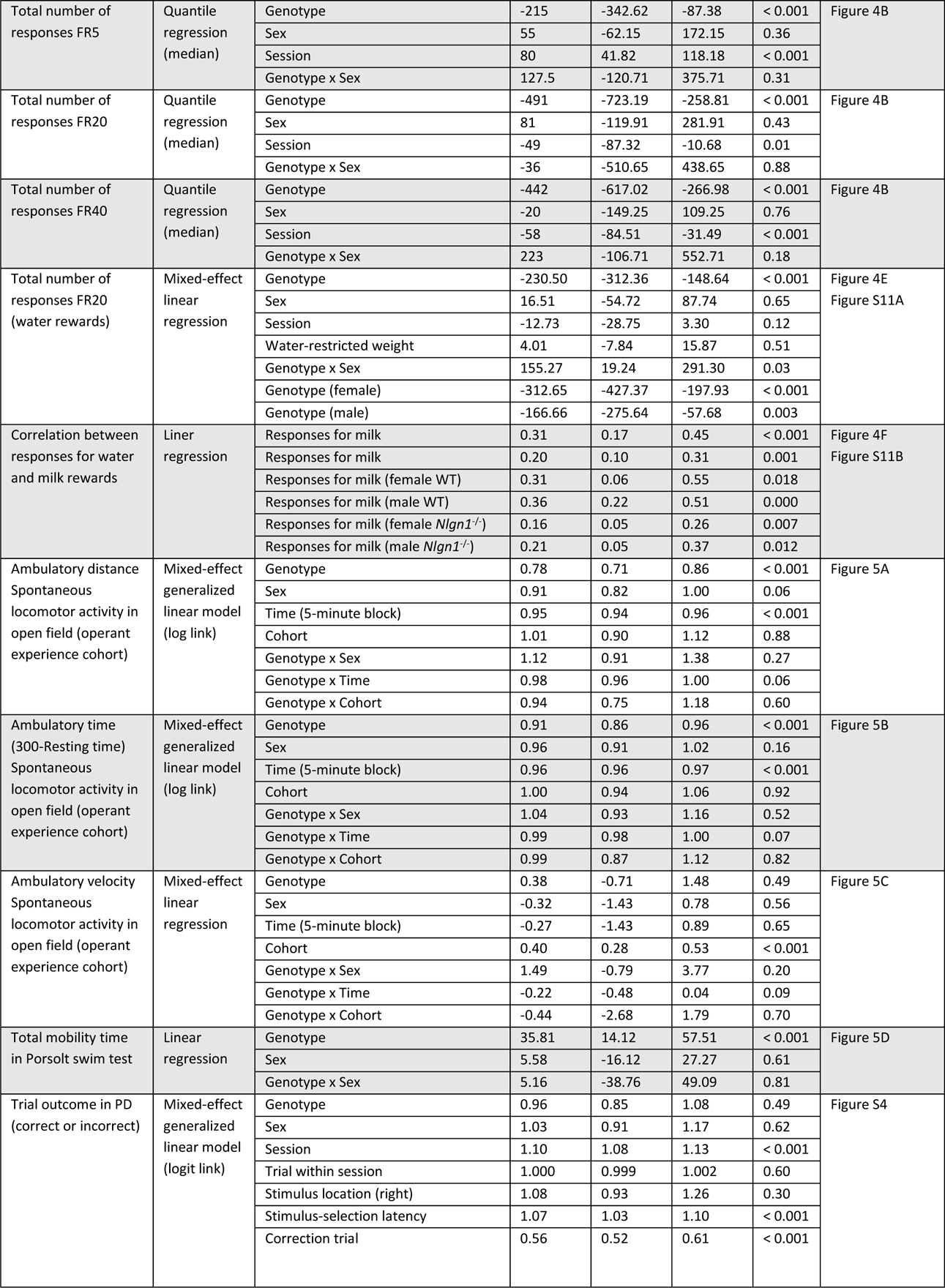
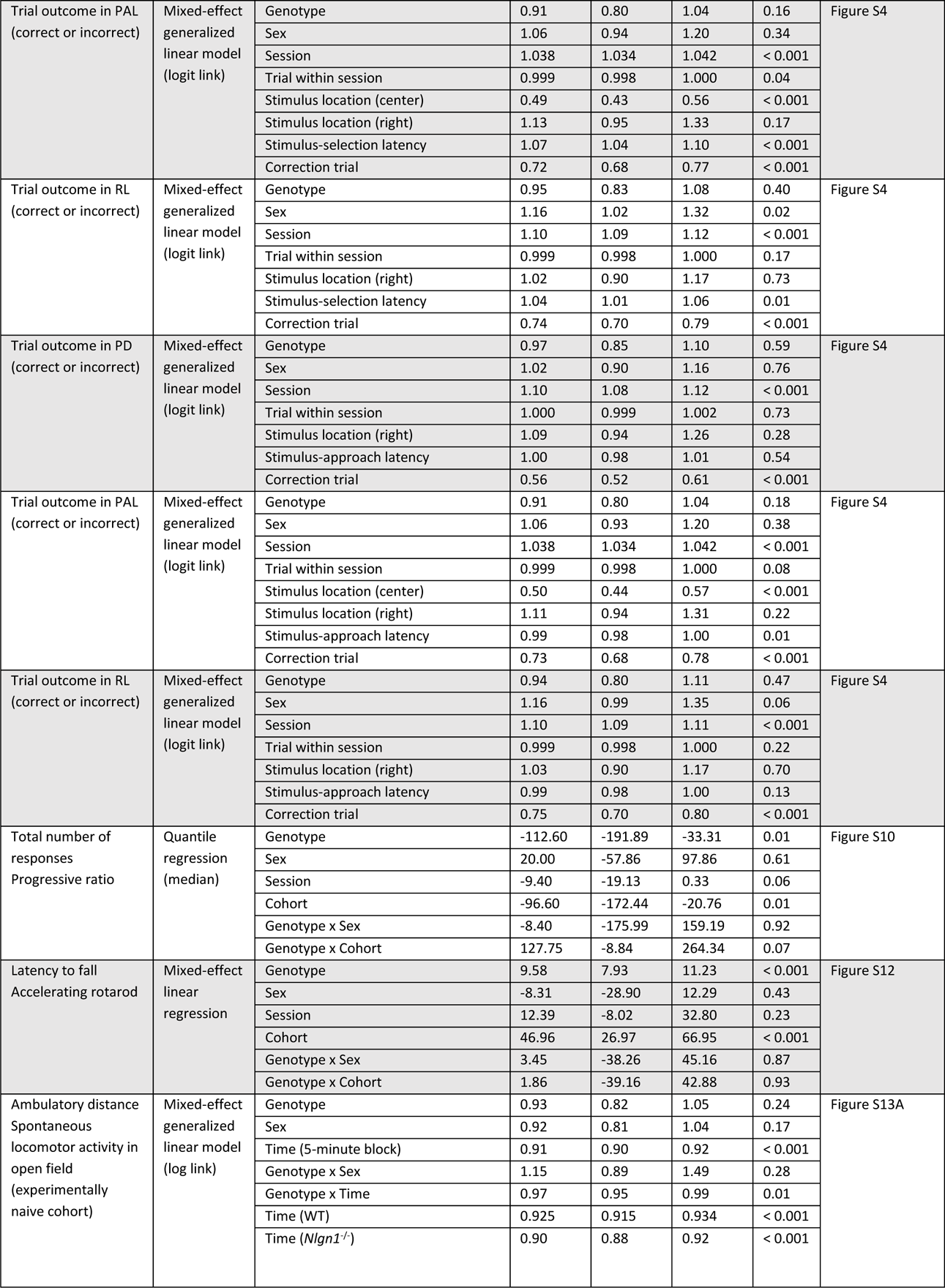
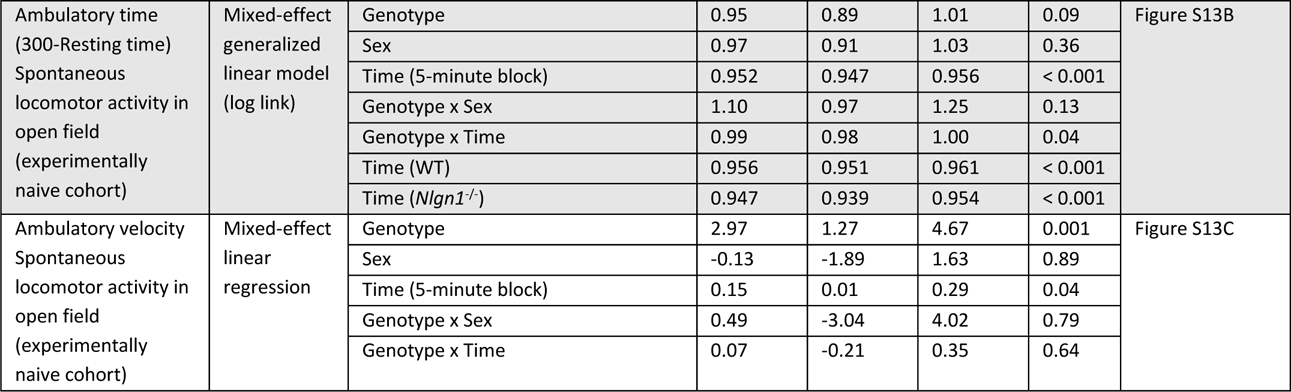
Variables included in regression models

## REFERENCES

1. Zador AM (2019) A critique of pure learning and what artificial neural networks can learn from animal brains. Nat Commun 10(1):3770.

2. Ichtchenko K, et al. (1995) Neuroligin 1: a splice site-specific ligand for beta-neurexins. Cell 81(3):435–443.

3. Song JY, Ichtchenko K, Sudhof TC, & Brose N (1999) Neuroligin 1 is a postsynaptic cell-adhesion molecule of excitatory synapses. Proc Natl Acad Sci U S A 96(3):1100–1105.

4. Irie M, et al. (1997) Binding of neuroligins to PSD-95. Science 277(5331):1511–1515.

5. Iida J, Hirabayashi S, Sato Y, & Hata Y (2004) Synaptic scaffolding molecule is involved in the synaptic clustering of neuroligin. Molecular and cellular neurosciences 27(4):497–508.

6. Haas KT, et al. (2018) Pre-post synaptic alignment through neuroligin-1 tunes synaptic transmission efficiency. eLife 7.

7. Barrow SL, et al. (2009) Neuroligin1: a cell adhesion molecule that recruits PSD-95 and NMDA receptors by distinct mechanisms during synaptogenesis. Neural development 4:17.

8. Budreck EC, et al. (2013) Neuroligin-1 controls synaptic abundance of NMDA-type glutamate receptors through extracellular coupling. Proc Natl Acad Sci U S A 110(2):725–730.

9. Heine M, et al. (2008) Activity-independent and subunit-specific recruitment of functional AMPA receptors at neurexin/neuroligin contacts. P Natl Acad Sci USA 105(52):20947–20952.

10. Mondin M, et al. (2011) Neurexin-Neuroligin Adhesions Capture Surface-Diffusing AMPA Receptors through PSD-95 Scaffolds. Journal of Neuroscience 31(38):13500–13515.

11. Blundell J, et al. (2010) Neuroligin-1 Deletion Results in Impaired Spatial Memory and Increased Repetitive Behavior. Journal of Neuroscience 30(6):2115–2129.

12. Chubykin AA, et al. (2007) Activity-dependent validation of excitatory versus inhibitory synapses by neuroligin-1 versus neuroligin-2. Neuron 54(6):919–931.

13. Espinosa F, Xuan Z, Liu S, & Powell CM (2015) Neuroligin 1 modulates striatal glutamatergic neurotransmission in a pathway and NMDAR subunit-specific manner. Frontiers in synaptic neuroscience 7:11.

14. Jiang M, et al. (2017) Conditional ablation of neuroligin-1 in CA1 pyramidal neurons blocks LTP by a cell-autonomous NMDA receptor-independent mechanism. Mol Psychiatry 22(3):375–383.

15. Jung SY, et al. (2010) Input-specific synaptic plasticity in the amygdala is regulated by neuroligin-1 via postsynaptic NMDA receptors. Proc Natl Acad Sci U S A 107(10):4710–4715.

16. Kim J, et al. (2008) Neuroligin-1 is required for normal expression of LTP and associative fear memory in the amygdala of adult animals. Proc Natl Acad Sci U S A 105(26):9087–9092.

17. Wu XT, et al. (2019) Neuroligin-1 Signaling Controls LTP and NMDA Receptors by Distinct Molecular Pathways. Neuron 102(3):621-+.

18. Nam CI & Chen L (2005) Postsynaptic assembly induced by neurexin-neuroligin interaction and neurotransmitter. P Natl Acad Sci USA 102(17):6137–6142.

19. Futai K, et al. (2007) Retrograde modulation of presynaptic release probability through signaling mediated by PSD-95-neuroligin. Nature neuroscience 10(2):186–195.

20. Kwon HB, et al. (2012) Neuroligin-1-dependent competition regulates cortical synaptogenesis and synapse number. Nature neuroscience 15(12):1667–1674.

21. Hoy JL, et al. (2013) Neuroligin1 drives synaptic and behavioral maturation through intracellular interactions. Journal of Neuroscience 33(22):9364–9384.

22. Jedlicka P, et al. (2015) Neuroligin-1 regulates excitatory synaptic transmission, LTP and EPSP-spike coupling in the dentate gyrus in vivo. Brain Struct Funct 220(1):47–58.

23. Shipman SL & Nicoll RA (2012) A subtype-specific function for the extracellular domain of neuroligin 1 in hippocampal LTP. Neuron 76(2):309–316.

24. Dahlhaus R, et al. (2010) Overexpression of the cell adhesion protein neuroligin-1 induces learning deficits and impairs synaptic plasticity by altering the ratio of excitation to inhibition in the hippocampus. Hippocampus 20(2):305–322.

25. Whitlock JR, Heynen AJ, Shuler MG, & Bear MF (2006) Learning induces long-term potentiation in the hippocampus. Science 313(5790):1093–1097.

26. Rogan MT, Staubli UV, & LeDoux JE (1997) Fear conditioning induces associative long-term potentiation in the amygdala. Nature 390(6660):604–607.

27. Reynolds JN, Hyland BI, & Wickens JR (2001) A cellular mechanism of reward-related learning. Nature 413(6851):67–70.

28. Nicoll RA (2017) A Brief History of Long-Term Potentiation. Neuron 93(2):281–290.

29. Nabavi S, et al. (2014) Engineering a memory with LTD and LTP. Nature 511(7509):348–352.

30. Moser EI, Krobert KA, Moser MB, & Morris RG (1998) Impaired spatial learning after saturation of long-term potentiation. Science 281(5385):2038–2042.

31. Morris RG, Anderson E, Lynch GS, & Baudry M (1986) Selective impairment of learning and blockade of long-term potentiation by an N-methyl-D-aspartate receptor antagonist, AP5. Nature 319(6056):774–776.

32. Giese KP, Fedorov NB, Filipkowski RK, & Silva AJ (1998) Autophosphorylation at Thr286 of the alpha calcium-calmodulin kinase II in LTP and learning. Science 279(5352):870–873.

33. Brigman JL, et al. (2008) Impaired discrimination learning in mice lacking the NMDA receptor NR2A subunit. Learn Mem 15(2):50–54.

34. Horner AE, et al. (2013) The touchscreen operant platform for testing learning and memory in rats and mice. Nature protocols 8(10):1961–1984.

35. Nithianantharajah J, et al. (2013) Synaptic scaffold evolution generated components of vertebrate cognitive complexity. Nature neuroscience 16(1):16–24.

36. Rabe-Hesketh S, Skrondal A, & Pickles A (2005) Maximum likelihood estimation of limited and discrete dependent variable models with nested random effects. J Econometrics 128(2):301–323.

37. Kalueff AV, et al. (2016) Neurobiology of rodent self-grooming and its value for translational neuroscience. Nature reviews. Neuroscience 17(1):45–59.

38. Crawley JN (2007) Mouse behavioral assays relevant to the symptoms of autism. Brain Pathol 17(4):448–459.

39. Mar AC, et al. (2013) The touchscreen operant platform for assessing executive function in rats and mice. Nature protocols 8(10):1985–2005.

40. Aberman JE & Salamone JD (1999) Nucleus accumbens dopamine depletions make rats more sensitive to high ratio requirements but do not impair primary food reinforcement. Neuroscience 92(2):545–552.

41. Niv Y, Daw ND, Joel D, & Dayan P (2007) Tonic dopamine: opportunity costs and the control of response vigor. Psychopharmacology 191(3):507–520.

42. Collins AG & Frank MJ (2014) Opponent actor learning (OpAL): modeling interactive effects of striatal dopamine on reinforcement learning and choice incentive. Psychological review 121(3):337–366.

43. Pessiglione M (2014) Why don’t you make an effort? Computational dissection of motivation disorders. European Psychiatry 29(8):541–541.

44. Pessiglione M, Vinckier F, Bouret S, Daunizeau J, & Le Bouc R (2017) Why not try harder? Computational approach to motivation deficits in neuro-psychiatric diseases. Brain.

45. Salamone JD & Correa M (2012) The mysterious motivational functions of mesolimbic dopamine. Neuron 76(3):470–485.

46. Salamone JD, Yohn SE, Lopez-Cruz L, San Miguel N, & Correa M (2016) Activational and effort-related aspects of motivation: neural mechanisms and implications for psychopathology. Brain 139(Pt 5):1325–1347.

47. Molendijk ML & de Kloet ER (2015) Immobility in the forced swim test is adaptive and does not reflect depression. Psychoneuroendocrino 62:389–391.

48. West AP (1990) Neurobehavioral Studies of Forced Swimming - the Role of Learning and Memory in the Forced Swim Test. Prog Neuro-Psychoph 14(6):863–877.

49. de Kloet ER & Molendijk ML (2016) Coping with the Forced Swim Stressor: Towards Understanding an Adaptive Mechanism. Neural Plast.

50. McClure SM, Daw ND, & Montague PR (2003) A computational substrate for incentive salience. Trends in neurosciences 26(8):423–428.

51. Dayan P (2012) Instrumental vigour in punishment and reward. European Journal of Neuroscience 35(7):1152–1168.

52. Brigman JL, et al. (2013) GluN2B in corticostriatal circuits governs choice learning and choice shifting. Nature neuroscience 16(8):1101–1110.

53. Ryan TJ, et al. (2013) Evolution of GluN2A/B cytoplasmic domains diversified vertebrate synaptic plasticity and behavior. Nature neuroscience 16(1):25–32.

54. Talpos JC, Winters BD, Dias R, Saksida LM, & Bussey TJ (2009) A novel touchscreen-automated paired-associate learning (PAL) task sensitive to pharmacological manipulation of the hippocampus: a translational rodent model of cognitive impairments in neurodegenerative disease. Psychopharmacology 205(1):157–168.

55. Marquardt K, et al. (2019) Impaired cognitive flexibility following NMDAR-GluN2B deletion is associated with altered orbitofrontal-striatal function. Neuroscience 404:338–352.

56. Skvortsova V, Palminteri S, & Pessiglione M (2014) Learning to minimize efforts versus maximizing rewards: computational principles and neural correlates. The Journal of neuroscience: the official journal of the Society for Neuroscience 34(47):15621–15630.

57. Prevost C, Pessiglione M, Metereau E, Clery-Melin ML, & Dreher JC (2010) Separate valuation subsystems for delay and effort decision costs. The Journal of neuroscience: the official journal of the Society for Neuroscience 30(42):14080–14090.

58. Samanez-Larkin GR, Hollon NG, Carstensen LL, & Knutson B (2008) Individual differences in insular sensitivity during loss anticipation predict avoidance learning. Psychological science 19(4):320–323.

59. Seymour B, et al. (2005) Opponent appetitive-aversive neural processes underlie predictive learning of pain relief. Nature neuroscience 8(9):1234–1240.

60. Pessiglione M, Seymour B, Flandin G, Dolan RJ, & Frith CD (2006) Dopamine-dependent prediction errors underpin reward-seeking behaviour in humans. Nature 442(7106):1042–1045.

61. Seymour B, Maruyama M, & De Martino B (2015) When is a loss a loss? Excitatory and inhibitory processes in loss-related decision-making. Curr Opin Behav Sci 5:122–127.

62. Pessiglione M & Delgado MR (2015) The good, the bad and the brain: neural correlates of appetitive and aversive values underlying decision making. Curr Opin Behav Sci 5:78–84.

63. Tye KM (2018) Neural Circuit Motifs in Valence Processing. Neuron 100(2):436–452.

64. Maia TV & Frank MJ (2011) From reinforcement learning models to psychiatric and neurological disorders. Nature neuroscience 14(2):154–162.

65. Brooks AM & Berns GS (2013) Aversive stimuli and loss in the mesocorticolimbic dopamine system. Trends in cognitive sciences 17(6):281–286.

66. Matsumoto M & Hikosaka O (2009) Two types of dopamine neuron distinctly convey positive and negative motivational signals. Nature 459(7248):837–841.

67. Hauser TU, Eldar E, & Dolan RJ (2017) Separate mesocortical and mesolimbic pathways encode effort and reward learning signals. P Natl Acad Sci USA 114(35):E7395–E7404.

68. Daw ND, Kakade S, & Dayan P (2002) Opponent interactions between serotonin and dopamine. Neural Networks 15(4-6):603–616.

69. Luo J, Norris RH, Gordon SL, & Nithianantharajah J (2018) Neurodevelopmental synaptopathies: Insights from behaviour in rodent models of synapse gene mutations. Prog Neuropsychopharmacol Biol Psychiatry 84(Pt B):424–439.

70. Varoqueaux F, et al. (2006) Neuroligins determine synapse maturation and function. Neuron 51(6):741–754.

71. Heath CJ, Bussey TJ, & Saksida LM (2015) Motivational assessment of mice using the touchscreen operant testing system: effects of dopaminergic drugs. Psychopharmacology 232(21-22):4043–4057.

72. Machado JAF, Parente PMDC, & Santos Silva JMC (2011) Qreg2: Stata module to perform quantile regression with robust and clustered standard errors. Statistical Software Components (Boston College Department of Economics), S457369.

73. Rescorla RA & Wagner AR (1972) A Theory of Pavlovian Conditioning: Variations in the Effectiveness of Reinforcement and Nonreinforcement. Classical Conditioning II: Current Research and Theory, eds Black AH & Prokasy WF (Appleton-Century-Crofts, New York), pp 64–99.

